# Attention improves information flow between neuronal populations without changing the communication subspace

**DOI:** 10.1101/2021.03.31.437940

**Authors:** Ramanujan Srinath, Douglas A. Ruff, Marlene R. Cohen

## Abstract

Visual attention allows observers to flexibly use or ignore visual information, suggesting that information can be flexibly routed between visual cortex and neurons involved in decision-making. We investigated the neural substrate of flexible information routing by analyzing the activity of populations of visual neurons in the medial temporal area (MT) and oculomotor neurons in the superior colliculus (SC) while rhesus monkeys switched spatial attention. We demonstrated that attention increases the efficacy of visuomotor communication: trial-to-trial variability of the population of SC neurons was better predicted by the activity of MT neurons (and vice versa) when attention was directed toward their joint receptive fields. Surprisingly, this improvement in prediction was not explained or accompanied by changes in the dimensionality of the shared subspace or in local or shared pairwise noise correlations. These results suggest a mechanism by which visual attention can affect perceptual decision-making without altering local neuronal representations.

## Introduction

Perhaps the most impressive hallmark of the nervous system is its flexibility. We effortlessly alternate between relying on or ignoring the same sensory information in different contexts. Visual attention dramatically affects perception and a wide variety of measures of neural activity in essentially every visual and visuomotor brain area (for reviews, see (Maunsell, 2015; Moore and Zirnsak, 2017)). Attention flexibly modulates signatures of neuronal activity including trial-averaged firing rates (Desimone and Duncan, 1995; Maunsell, 2015; Reynolds and Chelazzi, 2004), shared variability between pairs of neurons in the same (Cohen and Maunsell, 2009, 2011; Gregoriou et al., 2014; Herrero et al., 2013; Luo and Maunsell, 2015; Mayo and Maunsell, 2016; Mitchell et al., 2009; Nandy et al., 2017; Ni et al., 2018; Ruff and Cohen, 2014a, 2014b, 2016a, 2019; Verhoef and Maunsell, 2017; Yan et al., 2014; Zénon and Krauzlis, 2012) and different brain areas (Oemisch et al., 2015; Pooresmaeili et al., 2014; Ruff and Cohen, 2016a; Ruff et al., 2016), interdependence of neuronal populations on a range of timescales (Azouz and Gray, 2003; Bichot et al., 2005; Bosman et al., 2012; Briggs et al., 2013; Buffalo et al., 2011; Buschman and Miller, 2007; Dagnino et al., 2014; Fries, 2015; Fries et al., 2001; Gregoriou et al., 2009; Klink et al., 2017; Lakatos et al., 2008; Miller and Buschman, 2013; Moore and Armstrong, 2003; Ruff and Cohen, 2016a, 2017; Saalmann et al., 2007; Salinas and Sejnowski, 2001; Saproo and Serences, 2014; Womelsdorf and Fries, 2007; Womelsdorf et al., 2006a), and the dimensionality of population activity within each brain area (Cowley et al., 2020; Huang et al., 2019; Ruff et al., 2020).

The behavioral effects of attention make it clear that visual information can be flexibly routed: a stimulus can either guide or be unrelated to a perceptual decision depending on the task condition (Carrasco, 2011; Egeth and Yantis, 1997; Kohn et al., 2016a; Maunsell, 2015). In the visual system, neurons in each area send projections to a variety of different sensory, association, and motor areas, and only a small proportion of neuronal population activity is shared between even highly connected brain areas (Semedo et al., 2019). Recent work used correlative methods to identify a functional ‘communication subspace’, which consists of the dimensions of neuronal population space in which trial-to-trial variability is shared between areas (Semedo et al., 2019, 2020). We similarly adopt the term ‘communication’ to refer to functional communication (i.e., shared trial-to-trial variability in responses to the same visual stimulus).

An exciting possibility is that modulations in the shape or the constitution of this subspace could be a substrate for flexible, attention-dependent routing of sensory information. Compared to its behavioral effects, attention has remarkably modest effects on the amount of visual information encoded in visual cortex (Ruff and Cohen, 2019). Instantiating task or attentional flexibility via flexible routing rather than information coding could allow the brain to retain irrelevant visual information for future behavior or memory while the most relevant visual information guides behavior.

We investigated three potential mechanisms of flexible information flow between visual cortex and premotor neurons involved in decision-making. We tested the hypotheses that attention modulates information flow between areas by (1) changing the way visual or task information is represented in a local population, (2) changing the communication subspace itself, and/or (3) changing the efficacy of information transfer (Figure 1d).

**Figure 1:**
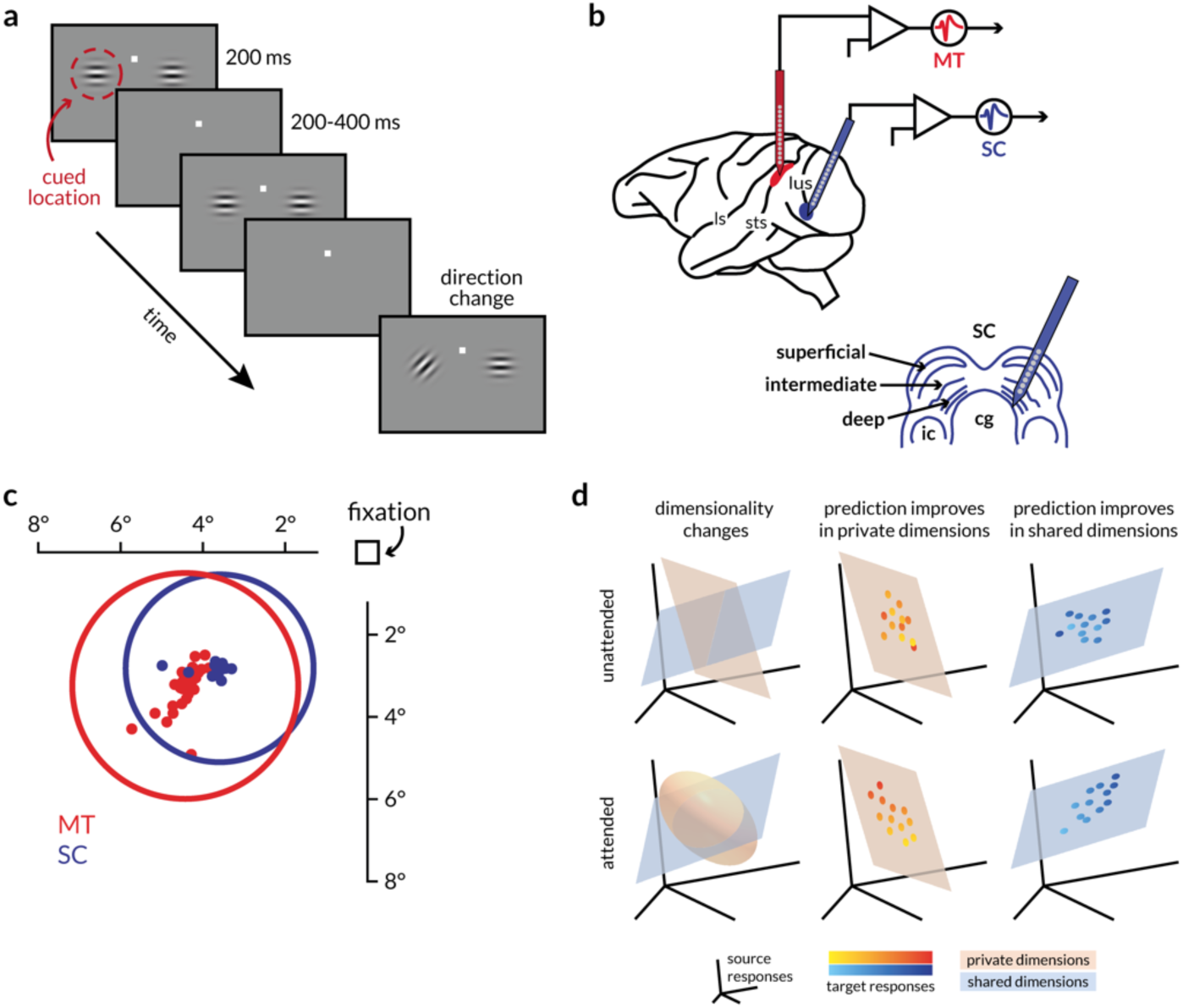
Behavioral task, recording sites, receptive fields, and schematic of hypotheses. **a:** Schematic of the motion direction change detection task. The monkeys were cued in blocks of trials to expect changes in motion direction at one of two spatial locations (cue was 80% valid). The monkey started the trial by fixating a central spot. Two small Gabor stimuli synchronously flashed on for 200ms and off for a randomized period of 200-400ms. One of the stimuli was positioned inside the joint receptive fields of the MT and SC neurons, and the other was placed in the opposite hemifield. Both stimuli moved in a direction that was chosen to drive the MT population well. After a randomized number of stimulus presentations (between 2 and 13), the direction of one of the stimuli changed. The monkeys were rewarded for making a saccade to the direction change in either location. We analyzed neuronal responses to all identical stimulus presentations except the first to minimize the effect of adaptation. **b:** Illustration of recording locations. Populations of MT and SC neurons were recorded with linear 24-channel moveable probes from the right hemisphere of two monkeys as they were doing the behavioral task described in (a). **c:** Receptive field locations of recorded units from an example recording session. The dots represent the receptive field centers of 28 MT (red) and 26 SC (blue) units. The circles represent the size and location of the median receptive field from each area. **d:** Schematics describing the hypotheses about attention-related changes in information flow between two areas. Each icon depicts the response space of the source area (the responses of the first n neurons or principal components, for instance), and orange and blue surfaces that represent two subspaces for the private or shared fluctuations in neural activity respectively. The two rows of icons represent the attended and unattended conditions (when attention was directed toward or away from the receptive fields of the recorded neurons), and each column describes the expected result of each of the following hypothesis. (left) Attention could alter the dimensionality of the private, shared, or both subspaces. If attention only modified local representations, then the number of private dimensions that explain the local neural fluctuations would change. (middle) Alternatively, attention could modulate information flow by enhancing or diminishing the extent to which neural activity in a target population tracks the neural activity of its source. If attention acted via this mechanism <u>locally</u>, then prediction would improve in <u>private</u> dimensions. (right) If attention modulated functional communication by modulating information flow <u>across areas</u>, then prediction would improve in <u>shared</u> dimensions.

Our strategy was to analyze functional communication between neuronal populations in visual and premotor areas while animals switched attention toward or away from their joint receptive fields. We recorded simultaneously from dozens of visual neurons in the medial temporal area (MT) and oculomotor neurons in the superior colliculus (SC) with overlapping receptive fields while rhesus monkeys performed a task in which they switched spatial attention, alternatingly using or ignoring the stimulus in the joint receptive fields of the recorded neurons. We used recently published methods for analyzing functional relationships between populations of neurons by assessing the dimensionality of shared variability and the extent to which the activity of one population could be predicted by the other (Semedo et al., 2019, 2020). We focused on trial-to-trial fluctuations in responses to the same visual stimulus because these are related to functional connectivity rather than simply reflecting tuning for similar stimuli (for review, see (Cohen and Kohn, 2011; Umakantha et al., 2020)), and have been shown to be correlated with choice behavior (Ni et al., 2018; Ruff et al., 2018).

We found strong evidence for our third hypothesis, that attention improves the efficacy of functional communication between visual and premotor neurons. Trial-to-trial variability of the population of SC neurons was better predicted by the activity of MT neurons (and vice versa) when attention was directed inside their joint receptive fields. This enhanced functional communication was not explained by increases or decreases in private or shared pairwise noise variability or a change in the number of private or shared dimensions of neuronal population activity.

This enhanced functional communication was not restricted to interactions between visual and premotor neurons. An independent dataset of simultaneously recorded neurons in primary visual cortex (V1) and in MT revealed that attention also increases functional communication within visual cortexEven though the attention-related change in pairwise correlations and response dimensionality within V1 was small compared to MT or SC, attention significantly enhanced our ability to predict the responses of single MT neurons from population activity in V1. Similarly, the effects of attention on functional communication were similar between MT and visual or motor neurons in the SC.

Our study provides a blueprint for combining behavioral paradigms that vary cognitive processes with dimensionality reduction and regression analyses to study how information can be flexibly routed in the nervous system. We used these methods to demonstrate that attention substantially improves the prediction performance between areas, more faithfully communicating information about attended stimuli, independent of changes in pairwise correlations or the dimensionality of either the local population or the shared variability. These results are the first demonstration of how attention affects the activity of distinct but connected populations of neurons in a way that affects the functional communication of visual information. They suggest a mechanism by which cognitive processes can affect perceptual decision making in ways that are independent of changes to the local neuronal representations.

## Results

We compared evidence consistent with several potential mechanisms for flexible routing of information. We chose a widely studied cued direction change detection task to study the behavioral effects of attention on visual perception, and three brain regions that are known to contribute to motion perception and visually-guided decision making – primary visual cortex (V1), the middle temporal area (MT), and the superior colliculus (SC). While rhesus monkeys performed the motion change detection task (Figure 1a), we recorded simultaneously from either dozens of neurons in MT and SC (Figure 1b) with overlapping receptive fields (Figure 1c; different aspects of these data were previously reported in Ruff and Cohen, 2019), or from several dozen neurons in V1 and a single MT neuron (Figure 6; different aspects of these data were previously reported in Ruff and Cohen, 2016a, 2016b). During the simultaneous MT-SC recordings, the monkey was cued as to which of two stimuli was most likely to change during a block of trials. This cued stimulus was placed either inside the overlapping receptive fields (RFs) of the recorded MT and SC neurons or in the opposite hemifield (Figure 1c). Throughout this manuscript, *attend in* refers to the trials where attention was directed toward the joint RFs and *attend out* refers to trials where attention was directed to the opposite hemifield. The monkey was rewarded for making a saccade to the location of the direction change, which occurred at a random and unsignaled time. The ability of the animal to detect the median difficulty changes in grating direction is enhanced by ∼ 25% on average across sessions when attention was directed to the location of the change (cued 76.5% detected, uncued 51.8% detected) (Ruff and Cohen, 2019). We analyzed the spike counts of each visually responsive multi-unit recorded from MT and SC during presentations of identical Gabor stimuli before the direction change (excluding the first presentation in each trial to remove adaptation effects). We also analyzed spike counts of each SC unit with elevated firing rates before saccade onset to the contralateral visual field. In the V1-MT data set, we tested our hypotheses on the responses of groups of V1 neurons whose receptive fields overlapped either of two small stimuli, both of which were inside the RF of the MT neuron (Ruff and Cohen, 2016a, 2016b).

### Signatures of population interactions that underlie attentional mechanisms

We tested the following non-mutually exclusive hypotheses (schematized in Figure 1d) about how attention might modulate information flow within and between areas. (a) Attention primarily modulates communication between areas by changing the dimensionality of either the private or the shared subspace (Figure 1d, left column). (b) Attention improves the fidelity of communication within local populations; this would be observable as an improvement in the ability to predict the activity of one subset of a neurons in a population from the activity of a different subset of neurons in the same area (Figure 1d, middle column) (c) Attention improves the fidelity of communication across brain regions; this would be evident in the improved accuracy of prediction of neural activity of one region using the activity of the other and vice versa (Figure 1d right column).

### Prediction of SC activity from MT activity using linear models improves with attention

Testing the predictions of our hypotheses requires calculating the ability to predict the activity of one population of neurons from another and identifying the dimensions of neural population space through which functional communication occurs. We plot the results of these analyses for one representative session in Figure 2. We used ridge regression to impose a sparse mapping between random subsets of MT neurons and the full populations of SC neurons in each attention condition (see Methods and Semedo et al, 2019).

**Figure 2:**
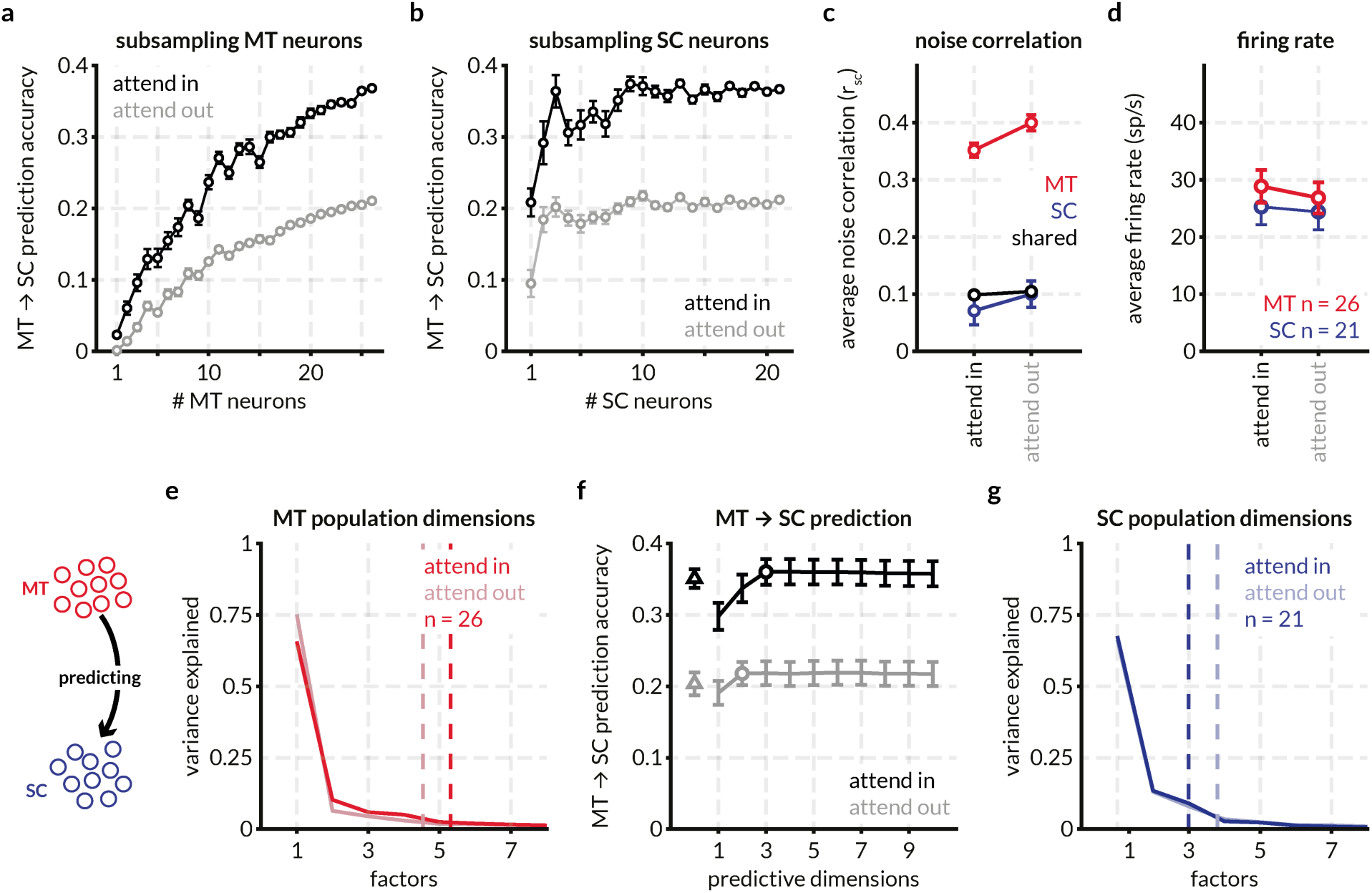
Attention improves prediction of SC activity from MT activity, increases firing rate, and decreases spike-count correlations in an example recording session. **a:** For an example session, the prediction accuracy of 1-26 randomly sampled (without replacement) MT neurons predicting the activity of a population of 21 SC neurons in the two attention conditions (attend in refers to the trials in which attention was directed within the joint RFs of the MT and SC neurons, and attend out refers to trials in which attention was directed in the opposite hemifield). Prediction was performed using a linear model with ridge regression and prediction performance was defined as the average cross-validated normalized square error (NSE) for the smallest ridge parameter for which the performance was within 1 SEM of the peak performance. Each point represents the mean prediction performance for n MT neurons predicting the full population of SC neurons. Error bars represent the standard error of the mean across random subsamples of n neurons. **b:** Same as (a) but for predicting random subsets of SC neurons using the activity of the full population of MT neurons, showing that the effect of attention on MT-SC communication is not limited to a subpopulation of the either the MT or SC neurons recorded in this session. **c:** Spike count correlation (r_SC_) defined as the correlation between the responses of pairs of neurons to all stimulus presentations for all MT neurons (325 pairs, red), SC neurons (210 pairs, blue), and MT-SC pairs (546 pairs, black). Attention decreases spike count correlations in MT (p = 1.2×10^-10^; Wilcoxon signed rank test (WSRT)) and SC (p = 0.0206; WSRT) but has no effect on pairwise correlations across areas (p = 0.2; WSRT) for this recording session. See Figure S1 for r_SC_ for all pairs across recording sessions. **d:** Neuronal firing rates increase with attention in MT (p = 8.3×10^-6^; WSRT) and SC (p = 0.04; WSRT) for this session. See Figure S1 for firing rates for all neurons across sessions. **e:** Factor analysis of MT population responses for this session reveals that 90% of the variance in the MT response fluctuations can be accounted for by ∼ 5 dimensions. The number of population dimensions is greater for the attend in condition vs the attend out condition. The arrow in the icon signifies that the MT population (source) is being used to predict the SC population (target): henceforth labeled as MT ➔ SC prediction. **f:** Predicting SC activity from MT responses using reduced-rank regression (RRR; black and gray lines) and ridge regression (triangle) reveals that the prediction performance for a matched number of trials is dramatically better for the attend in condition (black) vs the attend out condition (gray). The optimum number of dimensions (circle) for the reduced-rank regression was defined as the lowest number of dimensions for which prediction performance was within 1 SEM of peak performance. This performance is at least as good as the performance of the ridge regression performance that uses all the source dimensions for prediction (the difference between the RRR prediction and the ridge regression prediction was not significant across sessions; data not shown). The number of source dimensions required for optimum regression performance was 3 for attend in and 2 for attend out suggesting that fewer dimensions are required for communication between MT and SC than the total number of population dimensions. **g:** Factor analysis of SC neurons reveals that 90% of the variance in the SC response fluctuations can be accounted for by 3-4 dimensions. For this session, the number of population dimensions is greater for the attend out condition vs the attend in condition.

Several features of this example recording session were typical of our data set. First, no subset of the recorded MT neurons could effectively predict SC neural activity; the prediction accuracy monotonically increased with the addition of MT neurons. Second, the accuracy of prediction was significantly improved in the attend in trials vs attend out trials across all sub-selections of the MT population. Third, attention also improved the ability to predict random subsets of SC neurons from the full population of recorded MT neurons (Figure 2b).

To determine the relationship between these measures of functional communication between neuronal populations in MT and the SC and more well-studied effects of attention, we next calculated traditional metrics neuronal activity like pairwise spike count correlations (Figure 2c) and population firing rate (Figure 2d). For this session, attention significantly decreased spike count correlations in both MT and SC but did not have an effect on variability shared between pairs of neurons in different brain areas. Attention also significantly increased mean firing rates in this session. Firing rate and correlation changes across sessions are detailed in Figure S1.

For the example session, we observed no attention-related change in the population dimensionality in MT (∼ 5 dimensions; Figure 2e) and SC (∼ 3.5 dimensions; Figure 2g) defined as the smallest number of dimensions that captured 95% of the variance in the shared covariance matrix (assessed using factor analysis; (Cunningham and Yu, 2014); also see Methods for code and other resources).

We next tested whether, as between two areas of visual cortex (Semedo et al., 2019), interactions between MT and the SC are limited to a subset of dimensions of neural population space. For the example session in Figure 2, only 2-3 dimensions of MT activity (identified using reduced rank regression; see Methods; defined at the number of dimensions at which the curves in Figure 2f reach asymptote) predicted SC activity at least as well as a full linear model (fit using ridge regression; see Methods). The prediction accuracy for the attend in trials was significantly better than the attend out trials irrespective of the number of predictive dimensions (the black line is always above the gray line in Figure 2f).

### Attention improves prediction accuracy for inter-areal communication channels

Testing our hypothesized mechanisms of information flow (Figure 1) requires determining how attention affects the dimensionality and informativeness of interactions within and between populations of neurons in MT and the SC. We therefore fit linear models for repeated random splits of the populations of recorded MT and SC neurons in all four directions – MT ➔ SC, MT ➔ MT, SC ➔ MT, and SC ➔ SC. We depict the effect of attention on these four communication channels (for the same single session as in Figure 2) in the form of mean prediction accuracy across all tested population splits (Figure 3c-f). For this session, the prediction performance improves with attention for all functional communication channels except within MT where it depreciates. We estimated the population dimensionality as of each of the randomly split populations of MT and SC neurons using factor analysis to compare with number of predictive dimensions (Figures 3a, b). Consistent with result for the full population above, the number of dimensions within each area is not affected by attention.

**Figure 3:**
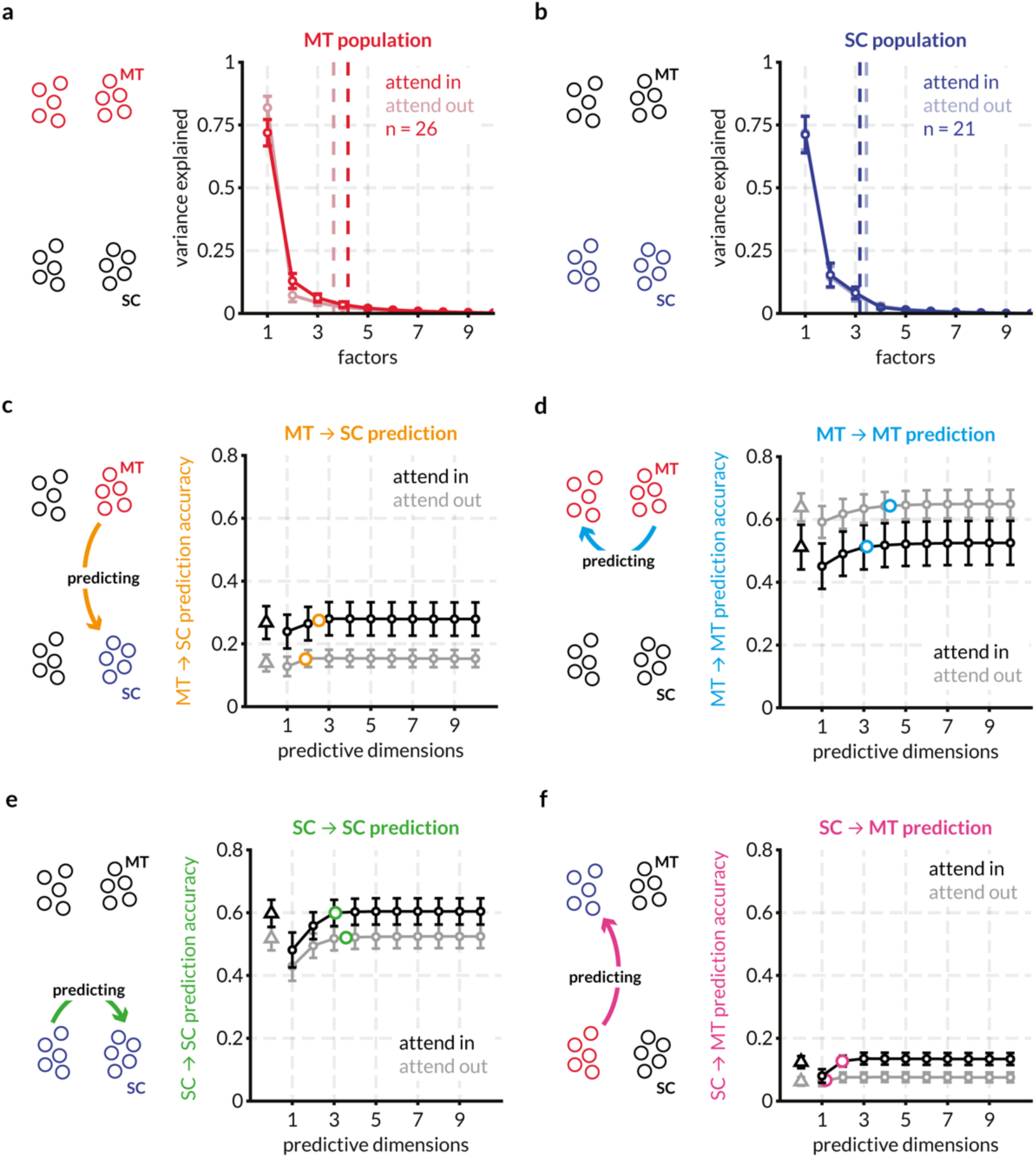
Randomly partitioned populations of MT and SC neurons predict activity within and across areas better with attention for the same example session. To compare prediction performance for inter-and intra-areal interactions, we randomly split both the populations of MT and SC neurons into two halves each – the target and source halves – as indicated in the icons. Each source half was used to predict the activity of both target halves using both the full linear model (ridge regression) and the reduced-rank regression (RRR) model. This split was done 20 times and the mean performance across the random splits is shown in c-f. Error bars indicate the SEM across these splits. **a:** Factor analysis of MT neurons reveals that 95% of the variance in the MT response fluctuations can be accounted for by ∼ 4 dimensions on average across all splits for this session. The number of population dimensions is greater for the attend in condition vs the attend out condition. **b:** Same as (a) for SC neurons. For this session, SC population fluctuations are captured by ∼ 3 dimensions in both attention conditions. **c:** Average prediction performance for the full model (black and gray triangles) and the RRR model (black and grey circles) across random splits of the MT and SC populations. The orange circle indicates the average optimum performance and average number of optimum prediction dimensions across the random splits. For each session, this point of optimum performance is plotted in different comparisons in the following figures. For this session, attention improves MT ➔ SC prediction performance. For all predictions, the RRR model performs at least on par with the full linear model using ridge regression. **d:** same as (c) for MT ➔ MT predictions. For this session, attention degrades prediction performance. The average optimum performance and average optimum prediction dimensions are indicated with blue circles. **e:** same as (c) for SC ➔ SC predictions. For this session, attention improves prediction performance. The average optimum performance and average optimum prediction dimensions are indicated with green circles. **f:** same as (c) for SC ➔ MT predictions. For this session, attention improves prediction performance. The average optimum performance and average optimum prediction dimensions are indicated with pink circles.

Across sessions, prediction performance between MT and the SC improves with attention without changing the dimensionality of that communication (Figure 4). The number of predictive dimensions required to account for intra-areal communications was higher than the number of dimensions for inter areal communication in both attention conditions. Whereas prediction accuracy for intra-areal communication was consistently high and remained unaffected by attention, the prediction accuracy for inter-areal communication significantly improved with attention (Figure 4b, which shows the ratios of the number of predictive dimensions and of the prediction accuracy in the two attention conditions). Attention does not affect the number of predictive dimensions required for communication within and across areas (the marginal distributions of ratios are centered at and not significantly different from 1; Wilcoxon signed rank test) but improves the prediction accuracy between MT and the SC (the distributions of ratios of MT ➔ MT and SC ➔ SC prediction accuracy are centered at and not significantly different from 1 but the ratios of MT ➔ SC and SC ➔ MT prediction accuracy are significantly greater than 1; Wilcoxon signed rank test; see also the distributions for each communication channel in Figure S3).

**Figure 4:**
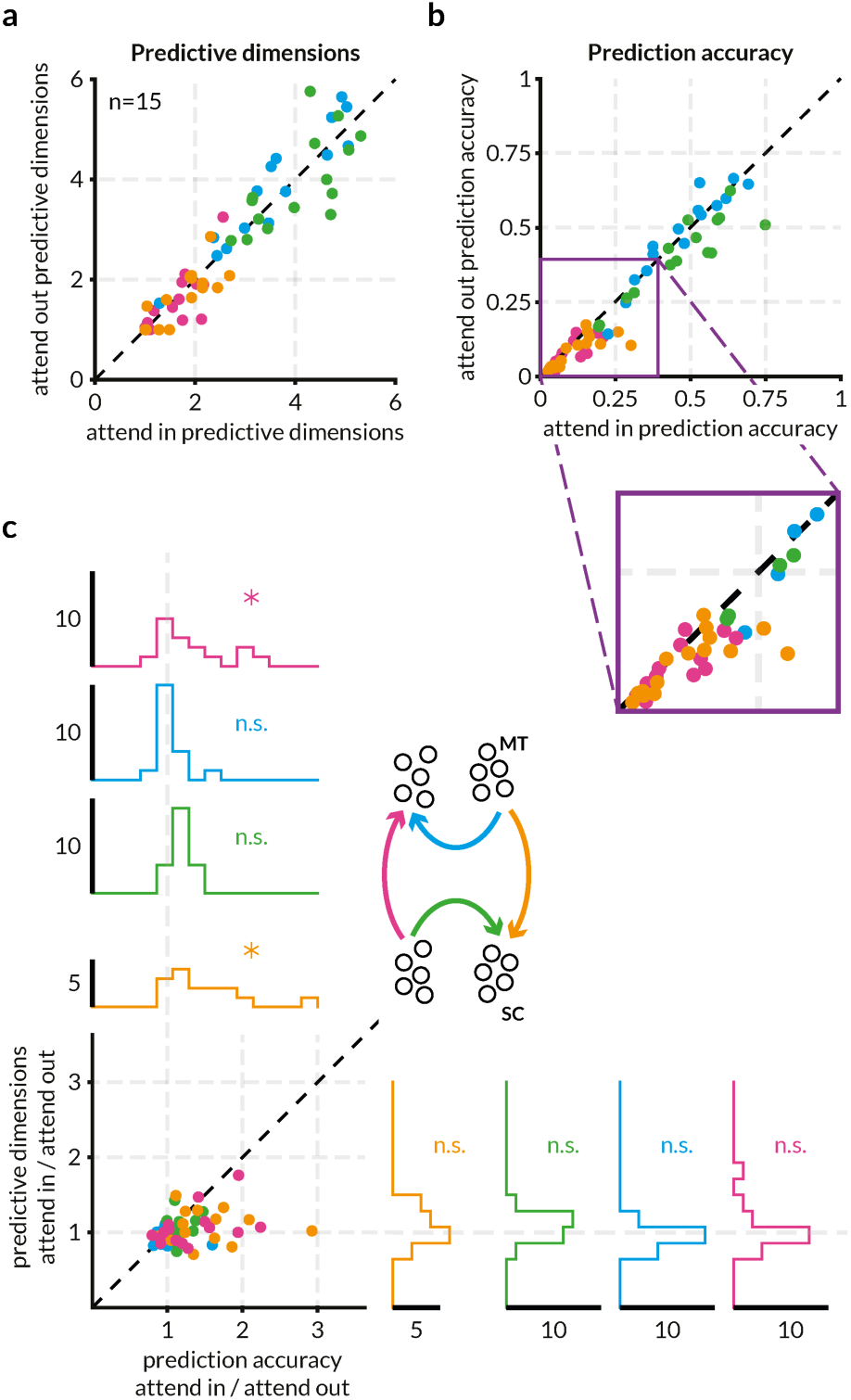
Attention improves the accuracy of across area prediction but not within area prediction without altering the dimensionality of the communication subspace. Each point of a given color represents a recording session. The color scheme is depicted in the icon in (c) and is consistent with other figures. **a:** Attention does not affect the dimensionality of the interaction between MT and SC neurons. Each point represents the average number of optimum predictive dimensions for each session for one of the four predictions – MT ➔ SC (orange), MT ➔ MT (blue), SC ➔ SC (green), SC ➔ MT (pink) – for the two attention conditions. There was so significant difference between the number of predictive dimensions for any of the four predictions. See Figure S5 for a detailed version of this panel. (MT-MT mean 3.67, range 1.5-5.2 for attend in and mean 3.74, range 1.1-5.3 for attend out; SC-SC mean 4, range 2.9-5.3 for attend in and mean 3.9, range 2.85-5.7 for attend out; MT-SC mean 1.8, range 1-2.5 for attend in and mean 1.75, range 1-2.7 for attend out; SC-MT mean 1.6, range 1-2.7 for attend in and mean 1.55, range 1-3.15 for attend out) **b:** Attention significantly increases the prediction accuracy of inter-areal but not intra-areal interactions. Each point represents the average prediction performance across random splits for one of the four predictions. The purple inset affords a zoomed in view of the relevant part of the plot which reveals that the points corresponding to the MT ➔ SC (orange) and SC ➔ MT (pink) predictions lie below the unity line. The average prediction accuracies for the attend in trials were significantly greater than those for the attend out trials for the MT ➔ SC prediction (p = 0.0015; Wilcoxon signed-rank test) and for the SC ➔ MT prediction (p = 8.54×10^-4^; Wilcoxon signed-rank test) but not the MT ➔ MT or SC ➔ SC predictions. **c:** The data in (a) and (b) visualized as a ratio of attend in and attend out. The marginal distributions of the ratios of prediction accuracy and predictive dimensions for all four predictions are also displayed. The mean ratios of prediction accuracy for MT ➔ SC (orange) and SC ➔ MT (pink) were significantly greater than 1 (p = 0.0016 and p = 0.012 respectively; t-test). The colored arrows in the icon indicate the source and target populations for each of the four predictions.

While attention is known to affect the mean pairwise spike count correlations within and between areas (Cohen and Maunsell, 2009; Mitchell et al., 2009; Ruff and Cohen, 2014a, 2016a), we found that attention-related improvements in prediction accuracy are not contingent on increases or decreases in spike count correlations. The ratio of prediction accuracies in the two attention conditions within and between areas was unrelated to the attention-related difference in mean spike count correlations between pairs of neurons within MT, within SC and between MT and SC (Figure S3).

The connectivity and functional roles of populations of SC neurons differ by layer, so we made use of our recordings that spanned layers to investigate whether functional communication between MT and the SC depends on layer as well. MT projections to SC predominantly end in the superficial layers in SC ((Fries, 1984, 1985) but also see (Lock et al., 2003)). Tecto-pulvinar projections from the superficial and intermediate layers of SC end in the inferior pulvinar which in turn projects to extra-striate areas (Lyon et al., 2010; Stepniewska et al., 1999). Also, there is some evidence that extra-striate projecting lateral geniculate nucleus (LGN) neurons do not receive direct retinal input and are dependent on SC projections across all layers for relaying visual information to MT (Benevento and Yoshida, 1981; Rodman et al., 1990). Given these laminar differences in cortical and thalamic inputs to and outputs from SC, we tested whether there is a difference between the attentional effect on information flow across functional classes of SC neurons. To classify SC neurons, we calculated an oculo-motor score based on SC neuron responses to visual stimuli and responses just prior to saccade onset (see Methods) and divided each population into two groups based on the rank ordering of oculo-motor scores. We then further split each SC population randomly as described before to serve as the source and target of regression with the simultaneously recorded MT population (Figure S4). We found no significant differences in the effect of attention on either the prediction accuracy or the number of dimensions required for prediction between the SC populations split by oculo-motor score (labeled *visual* and *motor* for brevity). Compared to random splits of the SC population, when split by oculo-motor score, the effect of attention on the prediction accuracy of the SC ➔ SC regression is pronounced (Figure S3c vs Figure S4c).

### Attention does not improve information flow by altering private or communication subspaces

The attention-related improvement in information flow (as implied by increased prediction accuracy across MT and SC) could in principle arise by changing the subspaces of activity responsible for functional communication within or between areas. We did not find evidence that attention changes the dimensionality of any of these subspaces: there was no attention-related change in the dimensionality of the local populations of MT and SC neurons (Figure S5a and S5b respectively) or in the number of predictive dimensions for the various communication subspaces within and between the two areas (Figure S5c-f). We consistently found that more dimensions were required to account for intra-areal communication than to account for inter-areal communication (mean 3.6 for MT ➔ MT and 4 for SC ➔ SC; vs 1.8 for MT ➔ SC and 1.6 for SC ➔ MT). This disparity suggests that MT and SC interact via a limited communication subspace.

Attention did not affect the dimensionality of the communication subspace. When we compared the number of dimensions used for private communication with the number of shared dimensions for MT ➔ MT prediction and MT ➔ SC prediction, we found that significantly fewer dimensions are required for MT ➔ SC communication than are available, but MT ➔ MT communication utilizes all available dimensions (Figure 5a). This effect was similar in the two attention conditions (Figures 5b and c), and we found similar results in the SC ➔ MT direction when compared with SC ➔ SC communication (Figure 5d-f). We found no relationship between the functional communication channels when assessed on a session-by-session basis (Figure S6). We also did cross-prediction analyses (using the attend in linear model to predict attend out data and vice versa) to check if the structure of the communication subspace changes while keeping its dimensionality, in turn causing the prediction accuracy to be better (Figure S7). We found that while the intra-areal models performed almost as well when swapped, the inter-areal models suffered a loss in prediction accuracy. This does not necessarily imply that the geometry of the communication subspaces changes with attention but that linear methods are unable to find a common subspace between the two attention conditions (also see Discussion).

**Figure 5:**
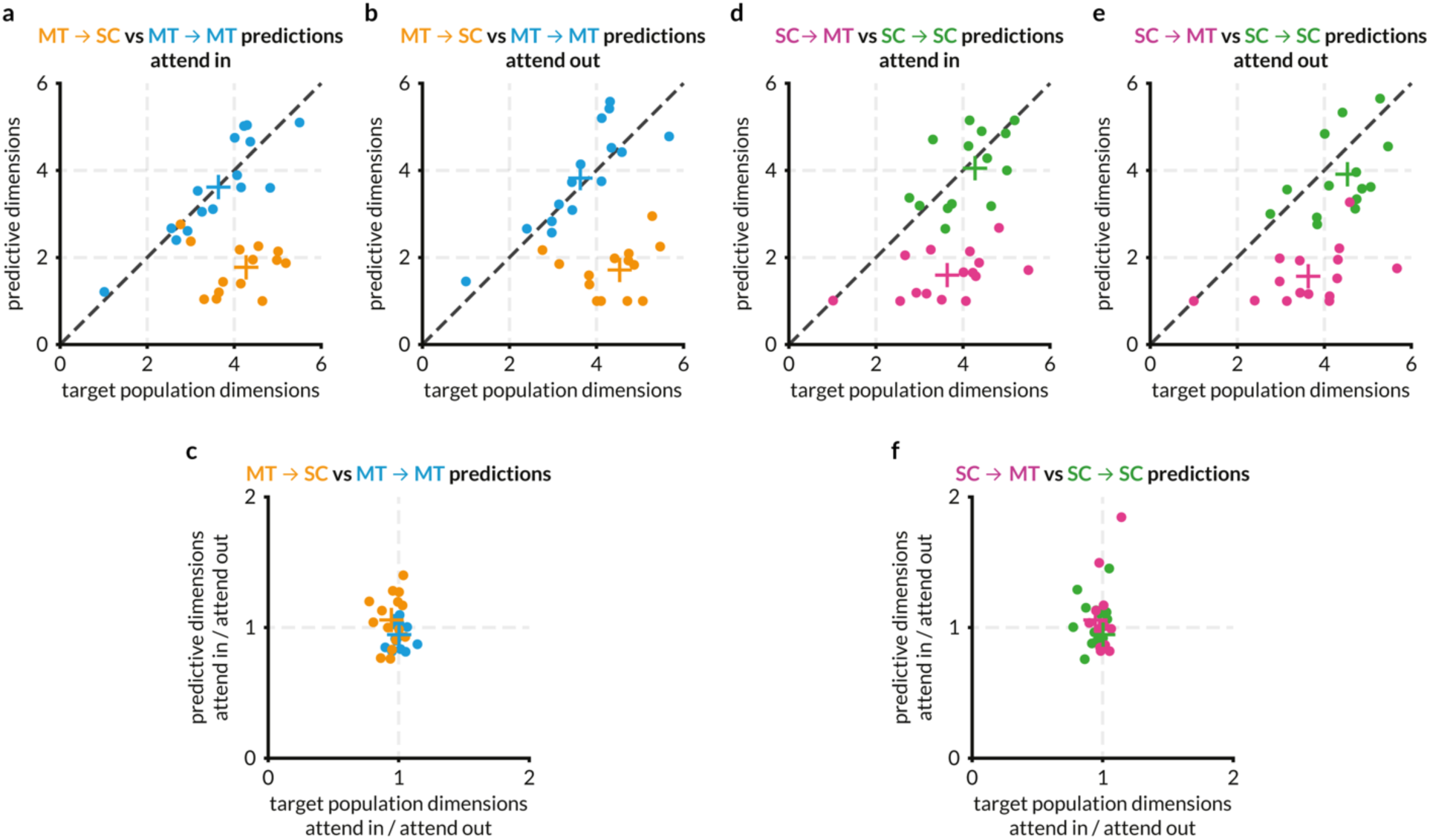
MT and SC populations interact via a communication subspace, but attention has no effect on the dimensionality of the communication subspace. Each point represents a recording session, and the color scheme is the same as other figures. Colored + represents the mean of the corresponding points. This figure compares the number of factors that explain 95% of the variance in the target area (from factor analysis) with the number of dimensions in the source area that are sufficient to predict the target area activity (from RR regression). Qualitative comparisons between the absolute values of the ‘number of dimensions’ from these two analyses in depicted in a, b, d, and e. The effect of attention is depicted in c and f. **a:** For the attend in condition, the number of private predictive dimensions are greater than the number of shared predictive dimensions in MT. Further, for the MT ➔ SC prediction (orange points), fewer dimensions are required to predict SC activity than are required to explain 95% of the variance in the SC activity, forming a communication subspace in MT that comprises of ∼ 2 shared dimensions that are sufficient to predict the ∼ 4-dimensional activity in SC. For the MT ➔ MT prediction, the number of predictive dimensions is similar to the number of population dimensions i.e., the predictive dimensions in MT are as large as possible and closely match the complexity of the target population, unlike the MT ➔ SC prediction. **b:** Same as (a) for the attend out condition. **c:** Data in (a) and (b) presented as a ratio to compare the effect of attention on the communication subspace in MT. **d:** For the attend in condition, the number of private predictive dimensions are greater than the number of shared predictive dimensions in SC. For the SC ➔ MT prediction (pink points), fewer dimensions are required to predict MT activity than are required to explain 95% of the variance in the MT activity i.e., a communication subspace exists in SC that comprises of ∼ 2 shared dimensions that are sufficient to predict the ∼ 3.5-dimensional activity in MT. **e:** Same as (d) for the attend out condition. **f:** Data in (d) and (e) presented as a ratio to compare the effect of attention on the communication subspace in SC.

### Attention improves information flow between V1 and MT

Both MT and SC exhibit relatively large attention-related changes in a number of measures of neuronal activity (Goldberg and Wurtz, 1972; Ignashchenkova et al., 2004; Krauzlis et al., 2013; Recanzone and Wurtz, 2000; Seidemann and Newsome, 1999; Womelsdorf et al., 2006b). Attention-related improvements in information flow may in principle be exclusive to pairs of regions that individually show significant changes in local representations. We tested this hypothesis by analyzing previously published simultaneous recordings of populations of neurons in V1 (which tend to show very modest effects of attention) and a single MT neuron (Ruff and Cohen, 2016a, 2016b; Ruff et al., 2016). As with the MT➔ SC results, we found that attention dramatically improves V1 ➔ MT prediction accuracy (Figure 6c; because we only recorded one MT neuron at a time, it was not possible to compute MT➔ V1 prediction accuracy). These results demonstrate that even though the effect of attention on V1 was small, attention-related effects on inter-areal communication are not contingent on large effects of attention in individual regions.

**Figure 6:**
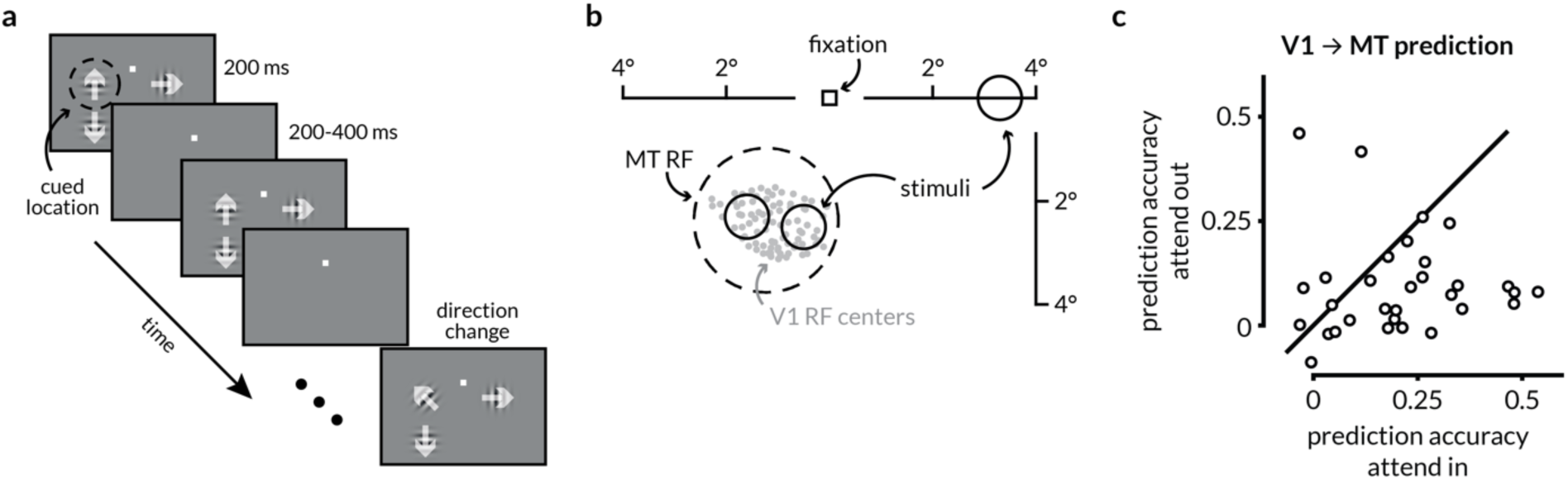
Attention enhances prediction accuracy between V1 and MT. **a:** Schematic of the motion direction change detection task used during the V1-MT recordings. The monkeys were instructed to attend to changes in motion direction at one of three spatial locations while ignoring changes at the other two locations in blocks of 50-100 trials. The monkey started the trial by fixating a central spot. Two or three small Gabor stimuli synchronously flashed on for 200ms and off for a randomized 200-400ms period. Two of the stimuli were positioned inside the joint receptive fields (RFs) of the V1 and MT neurons, and the other was in the opposite hemifield. Trials during which attention was directed into the MT RF towards either of the two spatial locations were considered attend in trials, and trials in which attention was directed to the opposite hemifield were considered attend out trials. In blocks when the monkey was cued to attend to one of the two locations inside the RFs, the third stimulus wasn’t presented. One of the two stimuli in the RF moved in the preferred direction of recorded MT neuron and the other moved in the anti-preferred direction. When presented, the third stimulus moved in the preferred direction of the MT neuron. After a randomized number of stimulus presentations (between 2 and 13), the direction of one of the stimuli changed. The monkeys were rewarded for making a saccade to the direction change in the cued location. Premature saccades or saccades to changes in motion direction at the un-cued location were not rewarded. We analyzed all identical stimulus presentations except the first to minimize the effect of adaptation. **b:** RF locations of recorded units from an example recording session. The gray dots represent the RF centers of 96 V1 neurons. The dotted circle represents the size and location of the RF for the recorded MT neuron. The size and locations of the stimuli were selected such that they lie within the MT RF. **c:** Attention improves the performance of V1 ➔ MT prediction. Each dot represents the cross-validated normalized r^2^ for a linear model of the MT neuron’s activity from V1 population activity using ridge regression for one recording session. The prediction accuracy on attend in trials was significantly greater than the accuracy on attend out trials (p = 0.0159; Wilcoxon signed-rank test). The value of the ridge parameter was chosen to be the smallest value for which the model performance was within 1 S.E.M. of the peak performance.

## Discussion

Our results show that attention changes the functional communication between populations of visual and premotor neurons. We demonstrated that attention changes the extent to which the activity of populations of neurons in the SC and be predicted by neuronal population in MT, and vice versa. These changes in information flow are not accompanied by changes in the dimensionality of the subspace of activity that is shared between areas, and they are independent of changes in firing rates, noise correlations, or population activity within each brain area. These results suggest that changes in information flow may mediate behavioral flexibility and place important constraints on models of flexible neural circuits.

### How attention-related increases in functional communication fit in with hypothesized mechanisms underlying attention

Previous studies have focused on a small number of hypothesized mechanisms by which attention might improve perception (Driver, 2001; Lavie, 2010; Peelen and Kastner, 2014; Ruff et al., 2018). The most studied hypothesis is that attention improves perception by improving information encoding (Cohen and Maunsell, 2009; Mitchell et al., 2007, 2009; Ruff and Cohen, 2014a). The observed attention-related changes in the responses of individual neurons and in correlations between visual neurons appear consistent with this hypothesis. However, neuronal populations typically encode more than enough sensory information to account for psychophysical performance (Kanitscheider et al., 2015; Kohn et al., 2016b; Parker and Newsome, 1998; Ruff and Cohen, 2014b, 2019), and the changes in trial-by-trial fluctuations may not reflect changes in information coding that are behaviorally-relevant (Baruni et al., 2015; Moreno-Bote et al., 2014). An alternate theory is that attention selectively improves the communication of sensory information to the neurons involved in perceptual decision-making. Physiological studies along these lines have primarily focused on changes in synchrony or coherence between areas on very short timescales (one or a few milliseconds, for review see (Womelsdorf and Fries, 2007)) or using human imaging data to assess functional connectivity over multiple seconds (Indovina and Macaluso, 2004; Ozaki, 2011; Rossi et al., 2014). However, co-variability on short timescales is mathematically nearly independent of correlations on the timescale of hundreds of milliseconds (Bair et al., 2001), and unlike fluctuations on very short or very long timescales, response fluctuations on the timescale of hundreds of milliseconds covary with perceptual decisions (Nienborg and Cumming, 2010; Nienborg et al., 2012).

Recently, we showed that attention is associated with only modest changes in either information coding in visual cortex or the way information is read out by premotor neurons on the timescale of perceptual decisions (Ruff and Cohen, 2019). Instead, our multi-neuron, multi-area recordings suggest that attention reshapes population activity in visual cortex which changes the visual information that guides behavior via relatively fixed readout mechanisms. Our current results suggest a functional implication of this reshaping, changing the information that is shared between sensory neurons and the premotor neurons involved in decision-making, without substantially changing the geometry of the subspace of activity that is shared between them.

### The communication subspace as a mechanism for flexible behavior

Many recent studies have shown that the activity of populations of neurons in many areas is generally confined to a subspace of population activity that is much lower dimensional than the number of recorded neurons (Cowley et al., 2016; Cunningham and Yu, 2014; Elsayed and Cunningham, 2017; Elsayed et al., 2016; Golub et al., 2016; Jazayeri and Afraz, 2017; Kaufman et al., 2014; Kiani et al., 2007; Miri et al., 2017; Morcos and Harvey, 2016; Pandarinath et al., 2018; Pitkow and Angelaki, 2017; Ruff et al., 2018; Sadtler et al., 2014; Yu et al., 2009). The divergent anatomical connections between even the most highly interconnected brain areas have long suggested that only a portion of the information encoded in each area is shared between areas.

A recent set of studies used recordings from multiple populations of neurons to establish that functional communication between different brain areas in the motor (Kaufman et al., 2014) or visual system (Semedo et al., 2019, 2021) is confined to a subspace of activity that is even lower dimensional than the activity within each area. Our results are consistent with the proposal in these that this limited communication subspace is an attractive mechanism for behavioral flexibility (Kaufman et al., 2014; Semedo et al., 2019). Because only a subset of information is shared, reshaping activity within the source area (as in Ruff and Cohen, 2019) and/or having a fixed but nonlinear subspace (proposed in Semedo et al., 2019) would change the information that is functionally communicated to a target area. Using cross-prediction analyses, we found that these linear methods reveal a difference in the structure of the communication subspace across attention conditions, but this observation may be consistent with a fixed, non-linear communication subspace, information flow could be improved by shifting the alignment of the shared fluctuations along the non-linearity (Figure S7). This mechanism is particularly attractive because changes in functional communication could occur without relying on changes in the weights relating one population to another, which may rely on synaptic plasticity mechanisms that occur over longer than behaviorally relevant timescales (Egeth and Yantis, 1997).

Our results demonstrate that the amount of information shared via the communication subspace between visual areas (V1 and MT, Figure 6) or between visual and premotor areas (MT and the SC, Figure 4) is in fact flexible. In future studies, it will be interesting to test the limits of this flexibility, such as whether this mechanism might mediate flexible communication of different stimulus features or information accumulated on different timescales that must mediate more complex forms of behavioral flexibility.

### Constraints on mechanistic models

Measurements of the activity of large populations has proven critical for constraining mechanistic models. Phenomenological models can explain attention-related changes in firing rates (Boynton, 2009; Ecker et al., 2016; Gilbert and Sigman, 2007; Maunsell, 2015; Navalpakkam and Itti, 2005; Reynolds and Heeger, 2009), but these do not provide insight into circuit mechanisms. A staggering variety of biophysical models can recreate the effects of attention on the trial-averaged responses of individual neurons (Ardid et al., 2007; Buia and Tiesinga, 2008; Deco and Thiele, 2011; Huang et al., 2019; Kanashiro et al., 2017; Silver, 2010; Sutherland et al., 2017). We and others have shown that attention-related changes in correlated variability that resides in a low dimensional subspace of population activity provides much stronger constraints on circuit models (Huang et al., 2019; Kanashiro et al., 2017).

The observations that functional communication between areas is lower dimensional than activity within each area (Kaufman et al., 2014; Semedo et al., 2020) and our observation that attention changes this communication will further constrain circuit models. In particular, many models (Brunel and Wang, 2001; Huang et al., 2019; Kanashiro et al., 2017; Machens et al., 2005; Rubin et al., 2015) and experiments (Fu et al., 2014; Karnani et al., 2016; Kuchibhotla et al., 2017) implicate inhibition in the flexibility of neuronal populations, but whether these mechanisms readily create low dimensional and flexible communication subspaces is unknown. It is possible that the complementary influence of different subtypes of inhibitory interneurons may underlie the flexible functional communication we observed (Cardin et al., 2009; Herrero et al., 2008; Roberts et al., 2005; Veit et al., 2017).

### Concluding remarks

The hallmark of the nervous system is its flexibility. Flexible behavior must rely, on some level, on flexible information flow. Attention, which changes the behavioral importance of different objects, features, or locations, is a good model of flexible information flow. Our results demonstrate that this flexibility is instantiated, at least in part, by changes in the information that is shared between different stages of the visuomotor pathway. These results lay the groundwork for establishing the role of flexible inter-area communications in a variety of sensory, cognitive, and motor computations.

## Acknowledgements

We are grateful to K. McKracken for providing technical assistance, to Adam Kohn for comments on an earlier version of this manuscript, and to Brent Doiron, Joao Semedo, and Byron Yu for helpful comments and suggestions regarding data analysis. M.R.C. is supported by National Institutes of Health grant R01EY022930 and Simons Foundation Grant 542961SPI.

## Author Contributions

Conceptualization, Methodology, Writing – Review & Editing, R.S., D.A.R., and M.R.C.; Software, R.S., D.A.R.; Analysis, Visualization, Writing – Original Draft, R.S. and M.R.C.; Funding Acquisition, Resources, Supervision, M.R.C.

## STAR ★ METHODS

### KEY RESOURCES TABLE

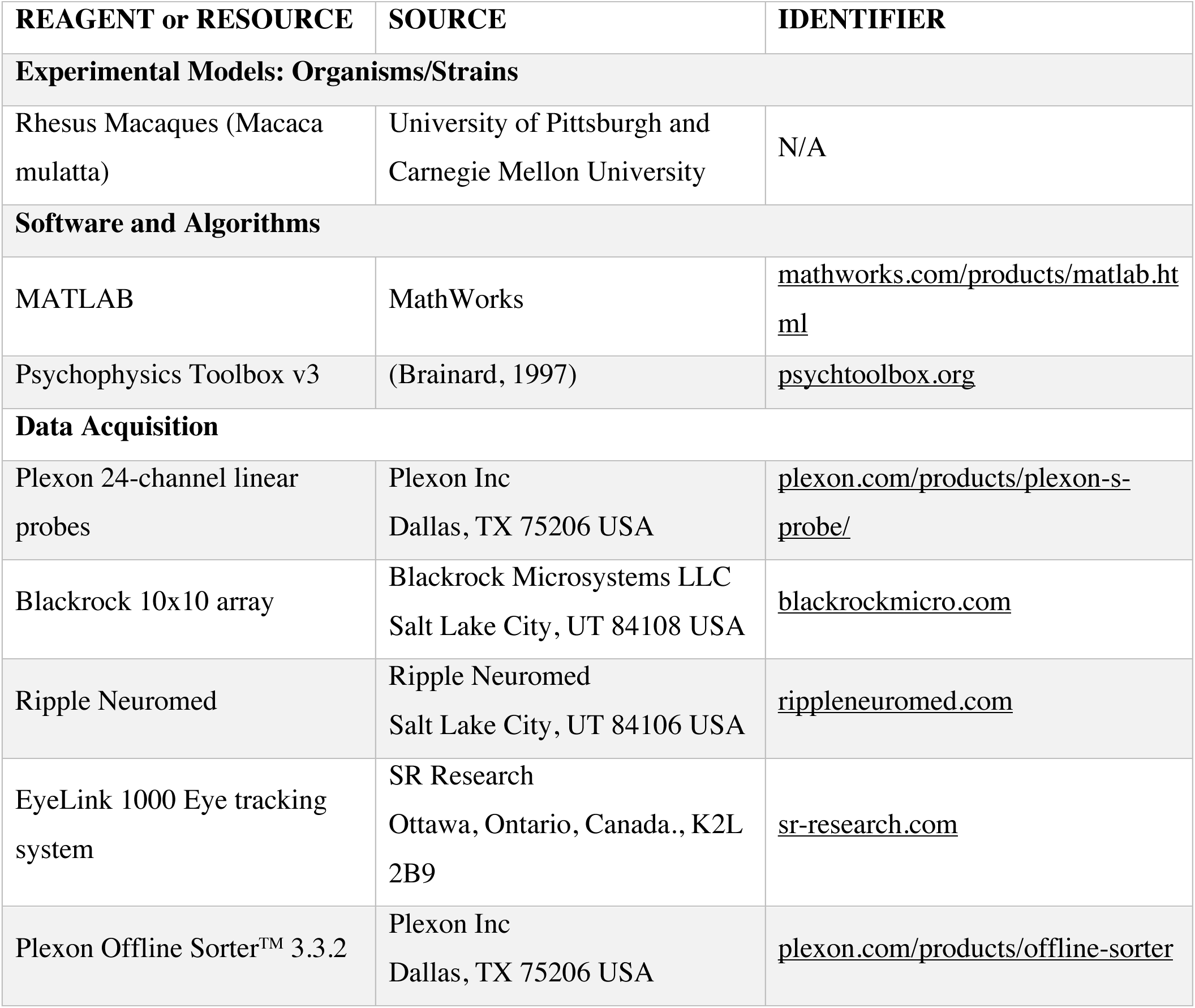

### RESOURCE AVAILABILITY

#### Lead Contact

Requests for resources should be directed to and will be fulfilled by the Lead Contact, Ramanujan Srinath.

#### Materials Availability

This study did not generate new unique reagents.

#### Data and Code Availability

The data and MATLAB code that support the findings of this study have been deposited in a public Github repository https://github.com/ramanujansrinath/mt-sc-comm-data. MATLAB code for reduced-rank regression and factor analysis has been publicly available by Byron Yu and can be downloaded from https://users.ece.cmu.edu/~byronyu/software.shtml. Further information and requests for data or custom MATLAB code should be directed to and will be fulfilled by the Lead Contact, Ramanujan Srinath.

### EXPERIMENTAL MODEL AND SUBJECT DETAILS

The electrophysiological data in this manuscript comes from two previously reported experiments (Ruff and Cohen, 2016a, 2019). In both experiments, two adult male rhesus monkeys (Macaca mulatta, 8 and 9 kg) were used. We implanted each animal with a titanium head post before behavioral training. We identified each cortical area by visualizing the sulci during array implantation, using stereotactic coordinates, and by observing the transition of grey and white matter signals on the movable probes. All animal procedures were approved by the Institutional Animal Care and Use Committees of the University of Pittsburgh and Carnegie Mellon University.

### METHOD DETAILS

#### Electrophysiological Recordings and Behavioral Task

Our methods for presenting visual stimuli and monitoring behavior have been described previously. Briefly, we presented visual stimuli using custom software (written in MATLAB using the Psychophysics Toolbox v3 (Brainard, 1997) on a cathode-ray tube monitor (calibrated to linearize intensity; 1,024 × 768 pixels; 120 Hz refresh rate) placed 54 cm from each animal. We monitored eye position using an infrared eye tracker (EyeLink 1000; SR Research) and recorded eye position and pupil diameter (1,000 samples/s), neuronal responses (30,000 samples/s) and the signal from a photodiode to align neuronal responses to stimulus presentation times (30,000 samples/s) using hardware from Ripple. All spike sorting was done offline manually using Offline Sorter (version 3.3.2; Plexon). We based our analyses on both single units and multiunit clusters and use the term *unit* to refer to either.

#### MT-SC recordings

We implanted two recording chambers on the right hemisphere that granted access to MT and SC for recordings with linear 24-channel moveable probes (Plexon; interelectrode spacing in MT = 50μm, SC = 100μm) and simultaneously recorded activity from neurons in MT and SC that had overlapping spatial receptive fields (Figure 1). To account for visual latencies in the two areas, spikes were counted between 50 and 250ms after stimulus onset. We only analyzed a recorded MT unit if its stimulus-driven firing rate was 10% higher than its firing rate as measured in the 100ms before the onset of the first stimulus. We only analyzed a recorded SC unit if its stimulus-driven firing rate was 10% higher than its firing rate as measured in the 100ms before the onset of the first stimulus or if its response during a 100ms epoch before a saccade on hit (correct) trials to the contralateral side was 10% larger than that same baseline. The dataset consisted of a total of 306 responsive MT units (6-29 units per session, mean 20.4) and 345 responsive SC units (12-29 units per session, mean 23) across 15 recording sessions. Each session began with receptive field mapping using a delayed-saccade task, and direction tuning during passive fixation, followed by multiple blocks of the following attention task. Each block began with a set of trials that instructed the monkey to attend to one of two spatial locations on the screen – either within the joint receptive fields of the neurons or in the opposite hemifield. Following that, each trial began when the monkey acquired fixation on a central spot within a 1.25° fixation window after which two peripheral drifting Gabor stimuli (one overlapping the receptive fields of the recorded neurons, the other in the opposite visual hemifield) synchronously flashed on (for 200ms) and off (for a randomized period between 200 and 400ms) between 3-12 times before, at a random, unsignaled time, the direction of one of the stimuli changed from that of the preceding stimulus. The monkey reported the orientation change by making a saccade to the changed stimulus within 450ms and received a juice reward. Each block consisted of approximately 100 completed trials (i.e., trials that ended in a hit or miss) after which the cued location of the orientation change switched to the other hemifield. Stimulus presentations during the response period of which the monkey made a micro-saccade were excluded from analysis. Neural responses to all stimulus presentations after the first (to minimize the effect of adaptation) and before the orientation change were analyzed. For each session, stimulus presentations were sampled such that the number of presentations was equal for each attention condition. Each session yielded 547-1909 (mean 1277) presentations for each attention condition. For each session, SC neurons were divided evenly into oculo-motor (*visual* for brevity) and motor neurons based on an oculo-motor score calculated as

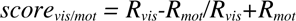

where *R_vis_* is the average neural response to the onset of the first stimulus, and *R_mot_* is the average response prior to a saccade to the target in the contralateral hemifield. This score was calculated for the trials where attention was directed into the joint RFs.

#### V1-MT recordings

We implanted a 10×10 chronic microelectrode array (Blackrock Microsystems) in V1 and a recording chamber to access MT. Each recording session began with searching a well-isolated MT neuron such that its receptive field (RF) overlapped the population RF of the V1 neurons and was driven similarly above baseline by a single stimulus flashed in each of two chosen locations. This dataset consisted of a total of 1631 responsive V1 units and 32 responsive MT units (1 unit per session in MT, 7–83 units per session, mean 51 in V1) across 32 recording sessions. Each block of trials began with a set of trials that instructed the monkey to attend to one of three spatial locations on the screen – either one of two locations within the receptive field of the MT neuron or one in the opposite hemifield. Each trial began when the monkey acquired fixation on a 1° fixation window. For blocks in which attention was directed within the RF of the MT neuron, two achromatic Gabor stimuli of equal contrast, spatial frequency, and speed were presented drifting in opposite directions (preferred and null direction for the MT neuron). For blocks in which attention was directed to the opposite hemifield, a third drifting Gabor was similarly flashed at the cued location. In these blocks, the contrast of the stimulus at the cued location was different from the two stimuli within the RF of the MT neuron. This was done to study the stimulus dependence of spike count correlations across cortical areas but is not critical to the current analyses as here the comparison is between the trials where attention is directed either into or out of the RF of the MT neuron regardless of stimulus parameters or specific location with the RF. After 2-14 presentations of the same stimuli, the direction of the stimulus at the cued location was changed and the monkey was rewarded for making a saccade to the changed stimulus within 500ms. As with the MT-SC data, stimulus presentations during which the monkey made a micro-saccade were excluded from analysis, all stimulus presentations after the first and before the orientation change were analyzed, and the presentations were sampled such that they were equal in the two attention conditions. Each session yielded 97-1469 (mean 583) presentations for each attention condition.

### QUANTIFICATION AND STATISTICAL ANALYSIS

#### Subsampling

To test whether attention affects prediction of neural responses within and across areas, we first sought to check whether or not the number of recorded neurons and trials across the two attention conditions in the datasets were sufficient for reasonable regression performance. We used a linear model of the form *Y=XB* where *X* and *Y* are matrices of *t x n* and *t x m* dimensions and *B* is the weight matrix of dimensions *n x m* (here, *t* is the number of stimulus presentations in a session, *m* and *n* are the numbers of neurons in the two areas). We found the ordinary least-squares solution for B by minimizing the squared prediction error as *B=(X^T^X)^-1^X^T^Y*. We sampled *N* MT neurons (where *N* went from 1 to the total number of recorded MT neurons) without replacement and used ridge regression predict SC responses. We did this subsampling 100 times for each *N*. For ridge regression, we chose the value of the regularization parameter (*λ*) using 10-fold cross-validation. The reported prediction accuracy corresponds to the largest *λ* for which mean performance (across folds) was within one SEM of the best performance. We also used the full MT recorded population to predict the responses of subsets of *N* SC neurons (where *N* went from 1 to the total number of recorded SC neurons) using the same method.

#### Noise correlations

The spike count correlation (r_SC_) was calculated as the correlation coefficient between the responses of the two units to repeated presentations of the same stimulus. Z-scoring responses before calculating noise correlations did not qualitatively change the comparisons between noise correlations and local or shared dimensionality or prediction accuracy. In Figure S1, noise correlations are computed for each pair in a session using all stimulus presentations in every trial (except the first), and then pooled across sessions and monkeys to yield 3315 pairs in MT, 3975 pairs in SC, and 6934 pairs across MT and SC. In Figure S3, noise correlations are computed as above and then averaged for each session.

#### Regression

To find the effect of attention on the ability of MT responses to predict SC responses and vice versa, we used the same linear model described above using ridge regression. This model is referred to as the full regression model in the text. To assess whether the SC activity can be predicted using a subset of MT population response dimensions (in other words, a subspace of MT activity), we used reduced-rank regression (RRR). The exact description and formulation of RRR can be found in (Semedo et al., 2019). Briefly, RRR constrains the weight matrix B to be of a given rank and is solved using singular value decomposition:

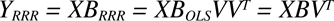

where *B_OLS_* is the coefficient matrix for the ordinary least-square solution, *B_RRR_* is the coefficient matrix for the RRR solution, *V* is a matrix whose columns contain the top principal components of the optimal linear predictor *Y_OLS_=XB_OLS_*. The columns of *B* define which dimensions of *X* are used for generating the prediction i.e., the predictive dimensions. As with the ridge regression solution above, we used 10-fold cross-validation and found the smallest number of dimensions for which predictive performance was within one SEM of the peak performance.

#### Cross-condition, cross-validated regression

To assess the effect of attention on the structure of the shared subspace between interaction populations of neurons, we calculated how well the regression weight matrix for one condition (attend in, say) predicted the responses of the target population in the opposite condition (attend out). In the first analysis, we simply used the cross-validated optimum number of dimensions to obtain a weight matrix in one condition and tested it against the trials of the other condition. The results of this method are depicted in Figure S7a-d. The accuracy of the inter-areal interaction dropped significantly but the accuracy of the intra-areal interaction was not affected. To assess whether this was a result of a linear scaling of the weight matrix across conditions due to non-stationarities or other task/stimulus independent factors, we projected the response of the source population using the weight matrix of the opposite condition before performing RR regression to obtain the prediction. This was cross-validated in the following way described in pseudo-code (for the MT ➔ SC interaction, for the attend out trials using the attend out vs attend in models, but we followed the same process for all potential permutations of conditions and areas).

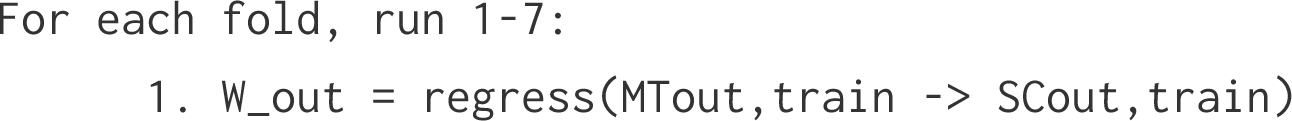
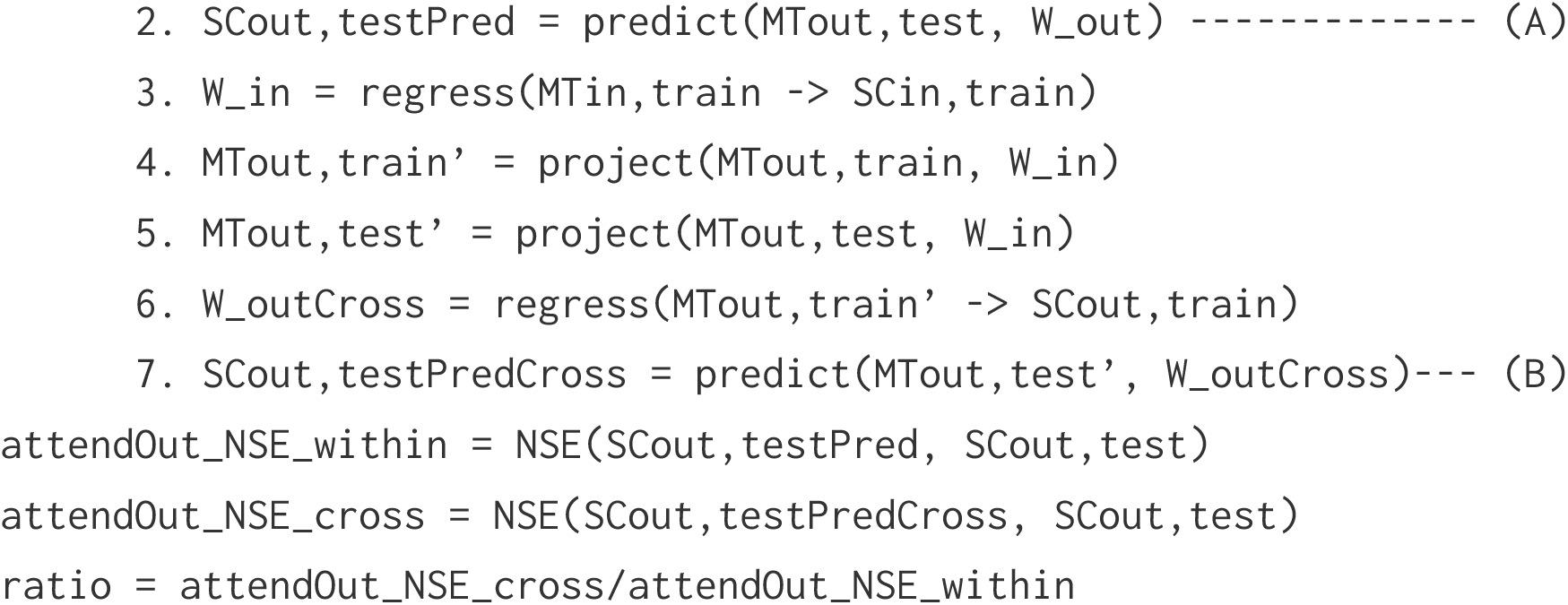

The ratio thus obtained was a cross-validated measure of how well the attend out weight matrix (`W_out`) performs compared to the weight matrix (`W_outCross`) that is trained to predict the same activity projected through the attend in weight matrix (`W_in`) first. We ran this for 10-folds for each random split of each population (described above) and evaluated the ratio of the normalized square error of prediction using both the matrices. This ratio is a quantitative measure of how well the cross-condition weight matrix performs relative to the within-condition weight matrix and values substantially lower than 1 would indicate a drastic drop in performance and, therefore, that the linear communication subspace between the two interacting populations is qualitatively different in their structure. We found this to be true for inter-areal interactions but not within-area interactions (Figure S7 e-h).

#### Factor Analysis

We used factor analysis (FA) to assess the dimensionality of neural activity within an area. FA is a static dimensionality reduction technique that does not assume the same noise variance for all recorded neurons and calculates the dimensions of greatest covariance (instead of variance). As with RRR, the details of this analysis can be found in previous publications (Everitt, 1984; Semedo et al., 2014; Yu et al., 2009). We followed the same steps as previously published work to estimate the dimensionality: (1) we found the number of dimensions *m_peak_* that maximized the cross-validated log-likelihood of the observed residuals; (2) we fitted a FA model with *m_peak_* dimensions and chose *m*, using the eigenvalue decomposition, as the smallest dimensionality that captured 95% of the variance in the shared covariance matrix. These population dimensions (*m*) and predictive dimensions as determined from RRR are determined by different techniques and therefore, wherever applicable, we have used these techniques to evaluate only the change of dimensionality (private or shared) between the two attention conditions instead of comparing absolute values.

## Supplementary Figures

**Figure S1 – related to figure 2:**
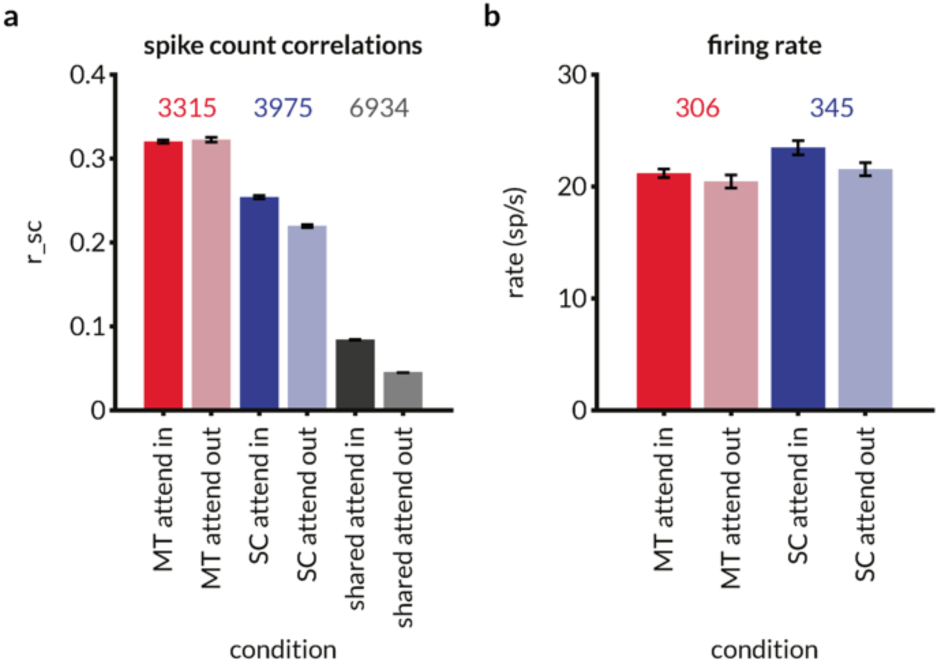
Effect of attention on aggregate noise correlations and firing rates for all neurons and pairs across all recording sessions. Error bars are standard error of the mean. **a:** Spike count correlations (r_SC_) for 3315 MT neuron pairs (red), 3975 SC neuron pairs (blue), and 6934 MT-SC pairs (gray) for attend in and attend out conditions. r_SC_ was calculated as the Pearson correlation between spike counts during all identical stimulus presentations except the first presentation after the beginning of the trial. Attention increases spike count correlations in SC pairs (p=2.7×10^-69^; Wilcoxon signed rank test) and MT-SC pairs (p=9.1 x10^-224^; Wilcoxon signed rank test) and has no effect on MT pairs (p=0.8; Wilcoxon signed rank test). The disparity between these results and previously published results is largely due to the selection of stimulus presentations. Here, we chose all presentations in a trial to increase statistical power in regression and factor analyses, whereas previous publications chose only the stimulus presentation before the orientation change to compare r_SC_ with behavioral outcomes. **b:** Average firing rate across all presentations for 306 MT neurons (red) and 345 SC neurons (blue). Attention significantly increases firing rates of neurons in both MT (p=8.87×10^-14^; Wilcoxon signed rank test) and SC (p=5.88×10^-42^; Wilcoxon signed rank test).

**Figure S2 – related to figure 4:**
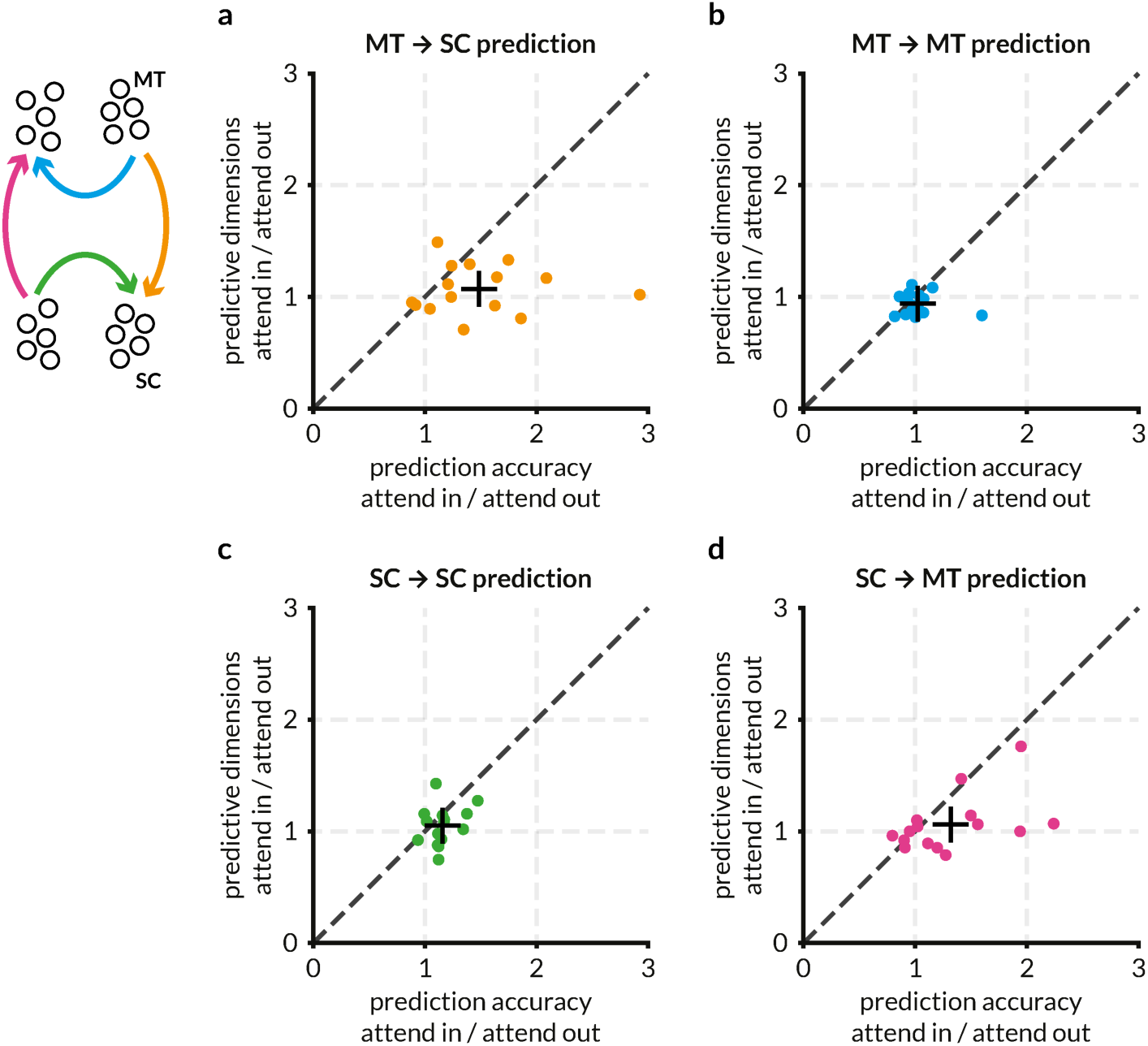
Attention improves prediction accuracy but not predictive dimensions for inter-areal communication. Each point represents a recording session, and the color scheme is the same as other figures and redundant with the plot labels. + represents the mean of the points. (same data as Figure 4c plotted separately for each prediction) **a:** Prediction accuracy and predictive dimensions presented as ratios between attend in and attend out conditions for the prediction of SC activity from MT activity. Each dot represents the average prediction accuracy and average predictive dimensions across 100 predictions of a random half of the SC population predicted by a random half of the MT population in that session. Attention increases prediction accuracy of MT ➔ SC predictions (p=0.0032; t-test) while having no effect on the number of predictive dimensions. **b:** Same as (a) but for MT ➔ MT predictions. Each dot represents the average prediction accuracy and average predictive dimensions across 100 predictions of a random half of the MT population predicted by the other half of the same population in that session. Attention has no effect on prediction accuracy or predictive dimensions. **c:** Same as (b) but for SC ➔ SC predictions. Attention has a small but significant effect on the prediction accuracy (p=7.9×10^-4^; t-test) but no effect on predictive dimensions. **d:** Same as (a) but for SC ➔ MT predictions. Attention increases prediction accuracy of SC ➔ MT predictions (p=0.0142; t-test) while having no effect on the number of predictive dimensions.

**Figure S3 – related to figure 4:**
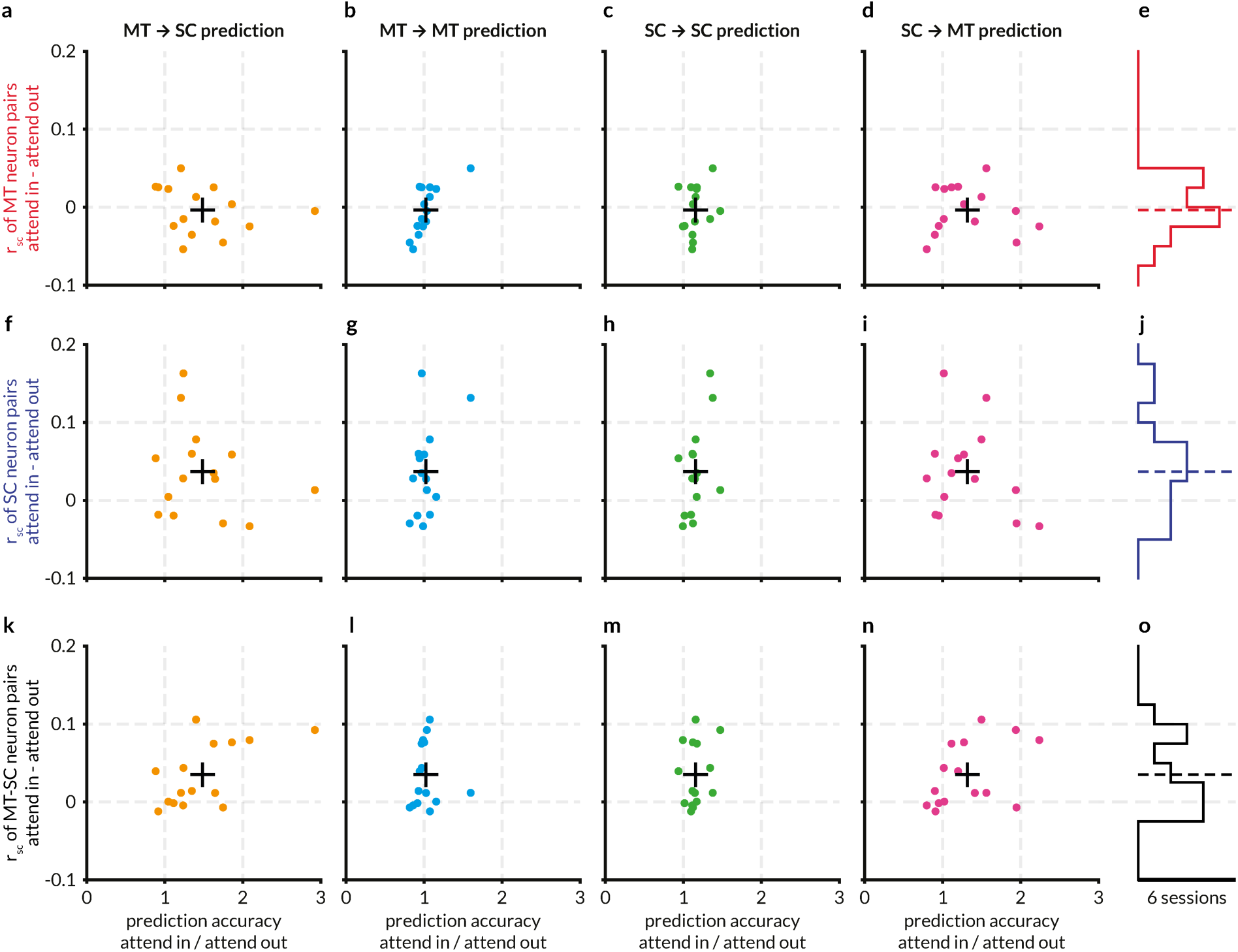
Attention-related changes in spike count correlations do not predict the improvement in communication efficacy across areas. Each panel illustrates how the differences of noise correlations of MT neuron pairs (a-e), SC neuron pairs (f-j), and MT-SC neuron pairs (k-o) between attend in and attend out conditions relate to the ratio of accuracies for within and across area response predictions. Each point represents a recording session, and the color scheme is the same as other figures and redundant with the plot labels. + represents the mean of the points. **a:** No relationship between the effect of attention on the average accuracy of MT ➔ SC predictions for each session and the effect on the average spike count correlations for MT neuron pairs for the same session. **b:** Same as (a) for MT ➔ MT predictions. **c:** Same as (a) for SC ➔ SC predictions. **d:** Same as (a) for SC ➔ MT predictions. **e:** Histogram of the difference of r_SC_ of MT neuron pairs between the two attention conditions. Dotted line represents the mean of −0.0035. **f-g:** Same as a-e, but for comparing prediction accuracies with session-wise average spike count correlations for SC neuron pairs. Dotted line in the histogram in (g) represents the mean of 0.0369. **k-o:** Same as a-e, but for comparing prediction accuracies with session-wise average spike count correlations for MT and SC neuron pairs. Dotted line in the histogram in (o) represents the mean of 0.0350. A weak relationship may be observed in (k) and (n) but the adjusted r^2^ for linear model fits are low (0.303 and 0.145 respectively) and not significant vs constant model.

**Figure S4 – related to figure 4:**
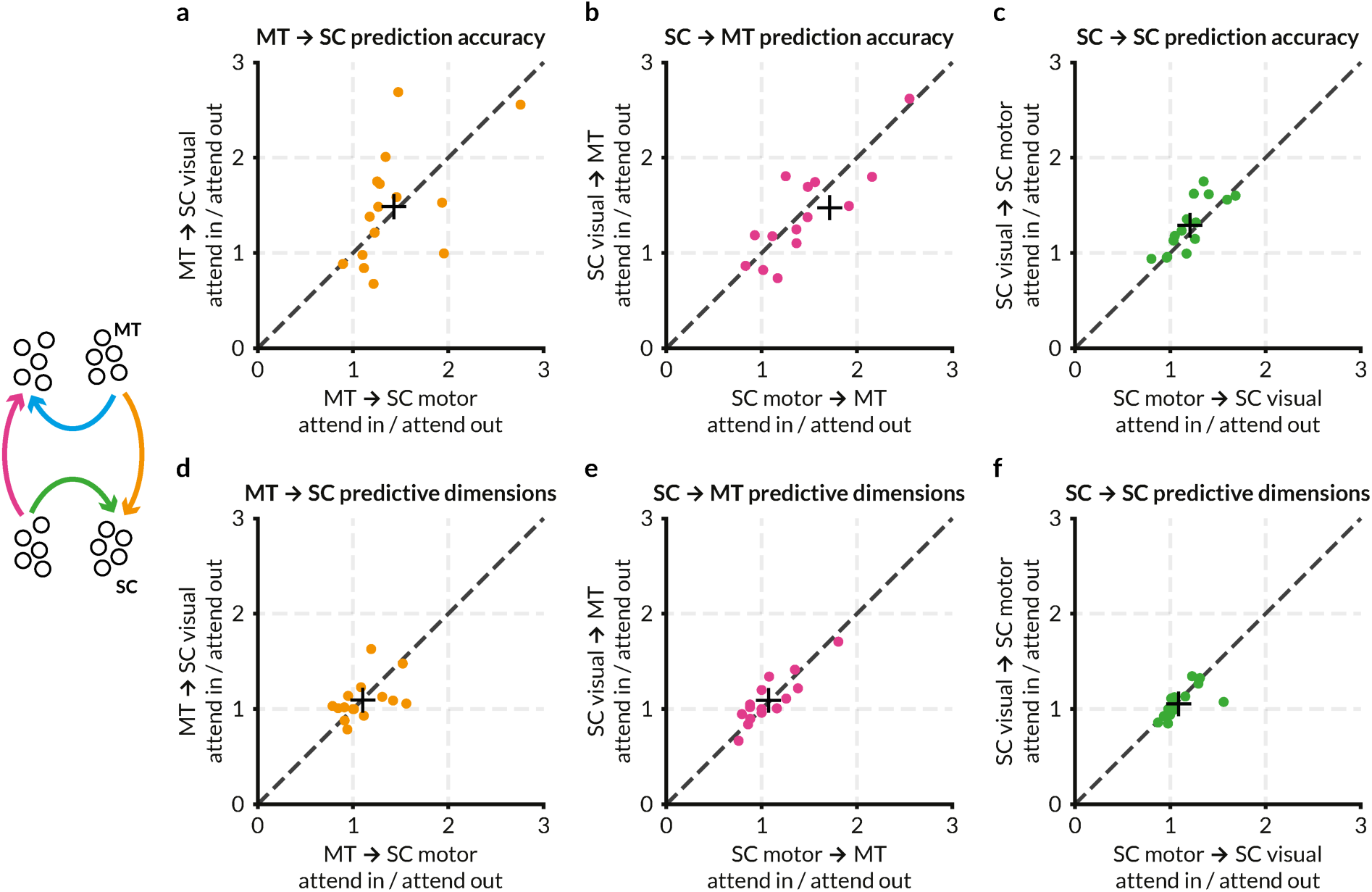
Both oculo-motor and motor neurons in SC contribute similarly to the attention-related improvement in prediction performance between MT and SC. For each session, SC neurons were ordered by an oculo-motor score (described in text and methods) and split evenly into “SC visual” and “SC motor” populations. (Oculo-motor SC neurons are labeled “SC visual” for brevity.) Each point represents a recording session, and the color scheme is the same as other figures and redundant with the plot labels. + represents the mean of the points. **a:** Average accuracy of predictions of randomly split SC populations of either oculo-motor neurons or motor neurons from the same population of randomly sampled MT populations presented as a ratio of the two attention conditions. (In each iteration, 50% of randomly sampled (without replacement) MT neurons were used to predict 50% of randomly sampled SC neurons from the top half of the oculo-motor index distribution and 50% of randomly sampled SC neurons from the bottom half of the distribution. So, effectively, only 25% of the SC neurons were used for predictions in these regressions as compared to 50% in other analyses.) The prediction accuracy of both oculo-motor SC and motor SC neural activity from MT neuron activity is similarly elevated with attention. Compare with figure 4c and supplementary figure 4a-a. (p = 0.0031 for MT ➔ SC motor, p = 0.0071 for MT ➔ SC visual, p = 0.52 for the ratio of the two; one-sample t-test for the ratios) **b:** Same as (a) for SC oculo-motor or motor ➔ MT predictions. As with (a), prediction accuracy is similarly enhanced with attention. Compare with figure 4c and supplementary figure 4a-d. (p = 0.0309 for SC motor ➔ MT, p = 0.0052 for SC visual ➔ MT, p = 0.456 for the ratio of the two; one-sample t-test for the ratios) **c:** Same as (a) for recurrent connections between SC oculo-motor and SC motor populations. As with (a), prediction accuracy is enhanced with attention. Compare with figure 4c and supplementary figure 4a-c. (p = 0.0047 for SC motor ➔ SC visual, p = 0.0013 for SC visual ➔ SC motor, p = 0.0495 for the ratio [SC visual ➔ SC motor] / [SC motor ➔ SC visual]) **d:** Same as (a) but for the ratio of the average number of predictive dimensions between the two attention conditions for the MT ➔ SC oculo-motor or SC motor predictions. Attention has no effect on the dimensionality of the shared subspace between MT and SC populations. Compare with figure 4c and supplementary figure 4a-a. (p > 0.05 for all ratios; t-test) **e:** Same as (b) for the ratio of the average number of predictive dimensions between the two attention conditions for the SC oculo-motor or SC motor predictions ➔ MT predictions. Compare with figure 4c and supplementary figure 4a-d. (p > 0.05 for all ratios; t-test) **f:** Same as (c) for the ratio of the average number of predictive dimensions between the two attention conditions for the recurrent connections between the SC oculo-motor and SC motor populations. Compare with figure 4c and supplementary figure 4a-c. (p > 0.05 for all ratios; t-test)

**Figure S5 – related to figure 5:**
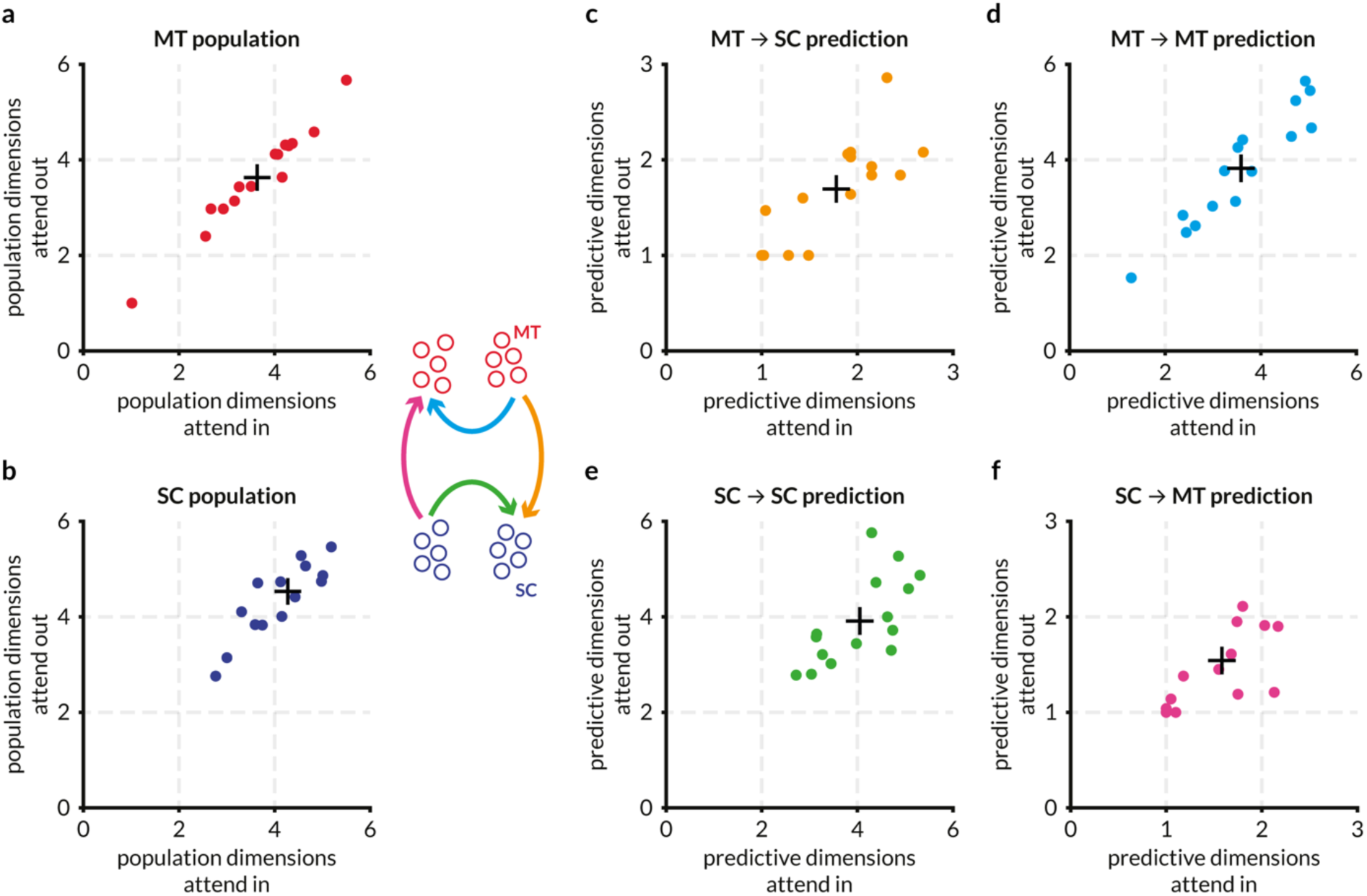
Attention does not alter the dimensionality of the response space in SC or MT, or the dimensionality of the shared communication subspace. Each point represents a recording session, and the color scheme is the same as other figures and redundant with the plot labels. + represents the mean of the points. **a:** Attention does not affect the population dimensionality of the MT populations. Each point represents the average number of dimensions (factors) required to explain 95% of the variance in the MT activity for one session. On average, fluctuations in MT activity are largely restricted to ∼ 3.5 dimensions. **b:** Attention does not affect the population dimensionality of the SC populations. Same as (a) for the SC population. On average, fluctuations in SC activity are largely restricted to ∼ 4.2 dimensions. **c-f:** Attention does not affect the number of dimensions required to optimally predict target activity for any of the four predictions. Same data as figure 4a split into four panels for clarity.

**Figure S6 – related to figure 5:**
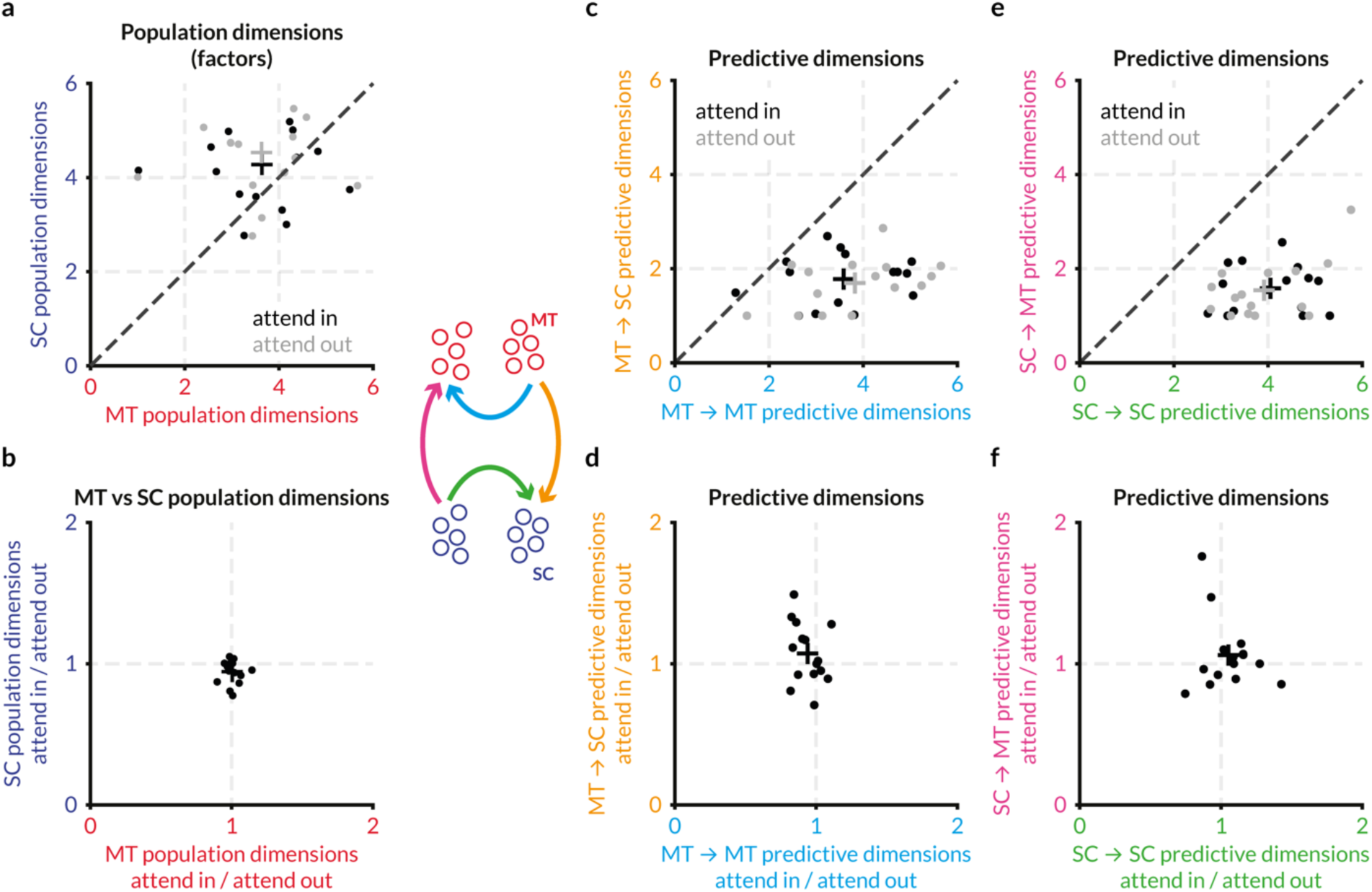
Detailed comparison of attention-related changes in MT and SC population dimensions and predictive dimensions different predictions. Each point represents a recording session, and the color scheme is the same as other figures and redundant with the plot labels. Colored + represents the mean of the corresponding points. **a:** Number of population dimensions or factors from factor analysis for the MT and SC populations in each session for attend in and attend out conditions. 95% of the variance in the MT and SC population activity can be explained with approximately 3.5 and 4.3 dimensions respectively in both attention conditions. **b:** Same as (a) expressed as a ratio of population dimensions in attend in and attend out conditions. Attention has no effect on the number of dimensions required to explain 95% of the variance in activity in this dataset. **c:** Number of predictive dimensions that are “shared” between MT and SC (orange axis) vs the number of dimensions that are “private” in MT (blue axis) in the two attention conditions. The number of MT dimensions required to predict SC activity (∼ 2) is lower than the number of MT dimensions required to predict MT activity (∼ 4). **d:** Same as (c) expressed as a ratio of predictive dimensions in attend in and attend out conditions. **e:** Same as (c) but for the number of dimensions in SC population activity that is sufficient to explain MT activity. Number of dimensions “shared” between SC and MT (∼ 2) in SC activity is lower than the number of “private” SC dimensions (∼ 4). **f:** Same as (e) expressed as a ratio of predictive dimensions in attend in and attend out conditions.

**Figure S7 – related to figure 5:**
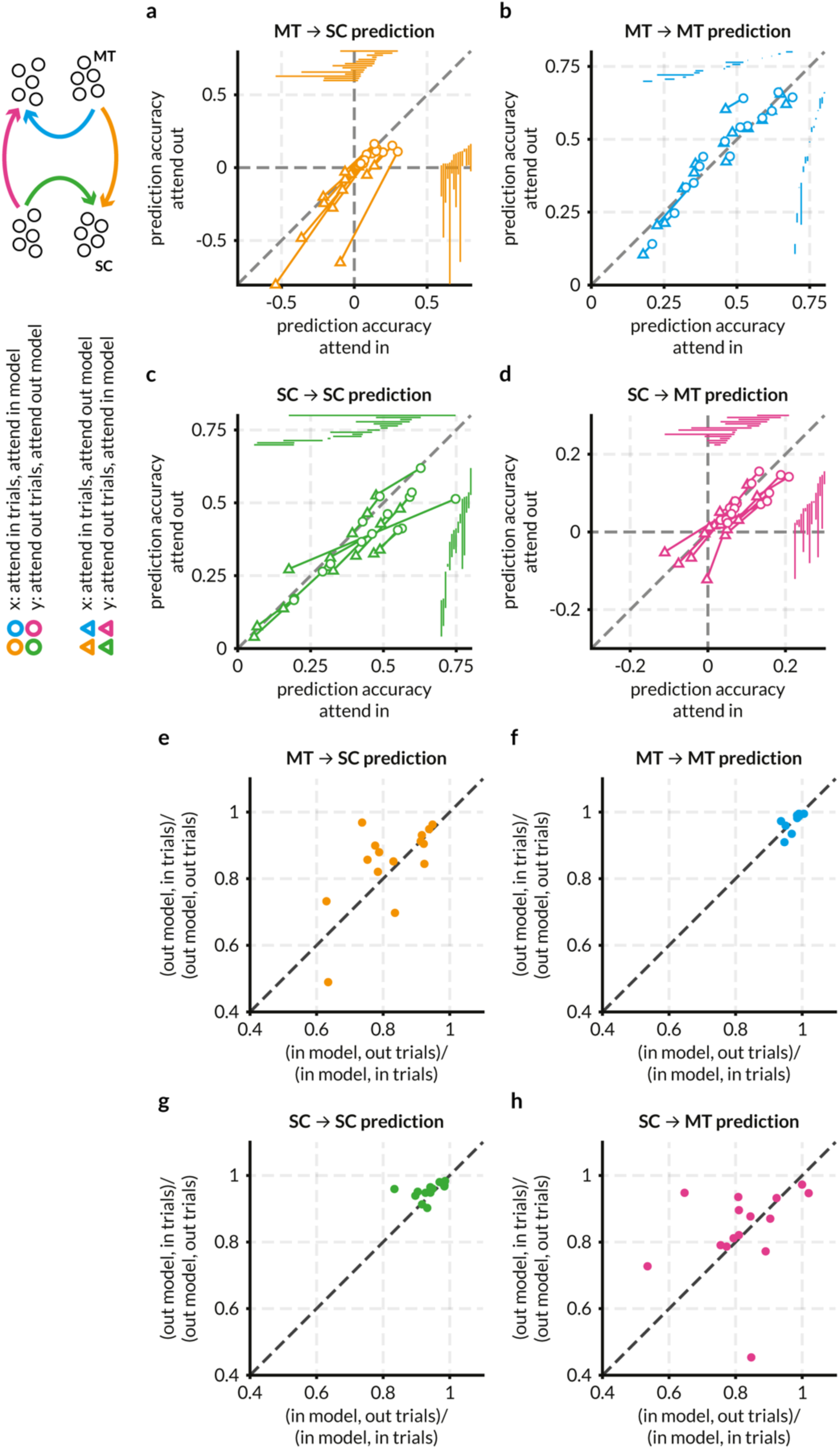
Cross-predicting activity for attend in trials using attend out model and vice versa reveals that the linear subspaces for across area communication are not identical. While the dimensionality of the communication subspace is not affected by attention, it is possible that the structure of the subspace changes while keeping its dimensionality, in turn causing the prediction accuracy to be better. To test this hypothesis, we used the weights of the linear model that corresponded to the optimum number of predictive dimensions in the attend in condition and used it to predict the target responses in the attend out condition and vice versa. We observed a marked drop in performance for cross-prediction for inter-areal communication in both directions but not intra-areal communication (a-d). To test whether this drop was due to a linear scaling of the weights across conditions and to cross-validate the cross-predictions, we projected the source activity through the weight matrix of the opposite attention condition and then fit a linear model to the target activity (see Methods for the details of the algorithm) and plotted the cross-validated cross-prediction performance normalized by the cross-validated performance of the true model. We observed a reduction in performance for the inter-areal predictions, albeit milder than earlier estimates (e-f). The intra-areal communication channels remained unaffected. While it may be possible that inter-areal communication indeed utilizes a different assortment of shared dimensions across attention conditions, we assert that these linear methods afford us a partial view of the effect of attention on the communication between areas. Each point represents the mean prediction accuracy of a recording session, and the color scheme is the same as other figures and redundant with the plot labels. **a:** We plotted the average cross-prediction accuracy (triangles) for each session and each communication channel across random splits against the true prediction accuracy (circles) i.e., the cross-validated prediction accuracy of the attend in models with the attend in trials etc. The linear model trained to predict SC activity using MT responses in the attend in condition performs significantly (p = 2.62×10^-4^; Wilcoxon rank sum test) worse when used to predict the SC responses for trials in the opposite attend out condition; the same is true for the reverse – using the attend out model to predict the attend in responses (p = 2.33×10^-5^; Wilcoxon rank sum test). Circles represent mean cross-validated prediction accuracy across random splits MT and SC neurons (same points as figure 4a). For each random split, the linear model of the opposite set of trials was used to predict the responses; the mean accuracy this out-of-set prediction across all random splits is represented by the triangles. Each circle-triangle pair is connected by a line and represents the change in prediction performance for a single session. The projections of each line on the cardinal axes are shown on the top and right of the plot, ordered by the prediction accuracy. Out-of-set prediction accuracies are always lower (p = 2.62×10^-4^; Wilcoxon rank sum test) and not significantly different from 0 (p = 0.07; t-test), which may mean that the model is unable to do better than guessing the target variance based on the mean of the target population activity (see Semedo et al., 2019 for more details). Both out-of-set models are similarly affected, evident from the consistent slope of the lines. This drastic drop in performance suggests that the shared communication subspace between MT and SC is different across attention conditions. **b:** Out-of-set mean accuracies for the MT ➔ MT prediction are not significantly different (p = 0.68 for the attend in model and p = 0.65 for the attend out model for attend in vs attend out trials; Wilcoxon rank sum test) suggesting not only that attention does not affect prediction performance within MT, but also that the same axes of fluctuations within the MT population activity are used for communication within MT thereby using the same private communication subspace. **c:** Same as (b) but for SC ➔ SC prediction. The out-of-set prediction is not significantly different (p = 0.046 for the attend in model and p = 0.097 for the attend out model for attend in vs attend out trials; Wilcoxon rank sum test). **d:** Same as (a) but for SC ➔ MT predictions. The out-of-set prediction is significantly worse for both the attend in model (p = 0.0011; Wilcoxon rank sum test) and the attend out model (p = 0.0016; Wilcoxon rank sum test). **e:** To control for the case where the prediction weights across conditions may be linearly scaled and thereby produce significantly worse predictions, the following procedure was used (these steps are for comparing the MT ➔ SC attend in weights with the attend out trials, but the same procedure applies for all possible permutations of conditions and populations). The pseudo-code for this cross-validated cross-prediction method can be found in Methods. First, the MT ➔ SC prediction weights were found for a set of attend out training trials (W_out) and the SC activity was predicted for the test trials (SCout,testPred). Similarly, the prediction weights for the training set of attend in trials was found (W_in). Then W_in was used to project the attend out MT activity for both training and test trials and then used to predict SC activity in the attend out condition for the test trials (SCout,testPredCross). After finding predictions across all folds, the normalized square error was found and compared for the within and across condition predictions. The ratio of the across/within condition prediction for the attend in trials for each session is plotted against the ratio of the across/within condition prediction for the attend out trials. This comparison between these variables demonstrates the ability of the same communication subspace being applied to the trials in the opposite condition and therefore a ratio substantially lower than 1 would indicate that the populations communicate using different subspaces in the different conditions. The cross-prediction accuracy is significantly lower for both attend in and attend out models tested with attend out and attend in trials respectively. **f:** same as **e**, but for MT ➔ MT interactions. As in **b**, the performance of the model from the opposite condition does not reduce prediction performance significantly. **g:** same as **e**, but for SC ➔ SC interactions. While the cross-prediction accuracy was not significantly different across the two attention conditions in **c**, the performance of the model was lower in each session. Here, the cross-validated cross-performance shows little difference in the ratio, which provides more evidence for the hypothesis that attention does not alter the dimensionality or structure of the SC-SC communication subspace. **h:** same as **e**, but for SC ➔ MT interactions. As in **e**, SC ➔ MT cross-prediction accuracy is significantly lower for both attend in and attend out models tested with attend out and attend in trials respectively. This difference in the structure or the constitution of the communication subspace between MT and SC between attention conditions may be evidence for attention either (a) altering the weights of interareal communication at a fast trial-to-trial timescale by unknown mechanisms, or (b) the inability of linear methods like FA and RRR to describe potentially non-linear response spaces and the non-linear dynamics of intra- and inter-areal interactions.

## References

Ardid, S., Wang, X.-J., and Compte, A. (2007). An Integrated Microcircuit Model of Attentional Processing in the Neocortex. J. Neurosci. 27, 8486–8495.

Azouz, R., and Gray, C.M. (2003). Adaptive Coincidence Detection and Dynamic Gain Control in Visual Cortical Neurons In Vivo. Neuron 37, 513–523.

Bair, W., Zohary, E., and Newsome, W.T. (2001). Correlated firing in macaque visual area MT: time scales and relationship to behavior. J. Neurosci. 21, 1676–1697.

Baruni, J.K., Lau, B., and Salzman, C.D. (2015). Reward expectation differentially modulates attentional behavior and activity in visual area V4. Nat. Neurosci. 18, 1656–1663.

Benevento, L.A., and Yoshida, K. (1981). The afferent and efferent organization of the lateral geniculo-prestriate pathways in the macaque monkey. J. Comp. Neurol. 203, 455–474.

Bichot, N.P., Rossi, A.F., and Desimone, R. (2005). Parallel and Serial Neural Mechanisms for Visual Search in Macaque Area V4. Science 308, 529–534.

Bosman, C.A., Schoffelen, J.-M., Brunet, N., Oostenveld, R., Bastos, A.M., Womelsdorf, T., Rubehn, B., Stieglitz, T., De Weerd, P., and Fries, P. (2012). Attentional stimulus selection through selective synchronization between monkey visual areas. Neuron 75, 875–888.

Boynton, G.M. (2009). A framework for describing the effects of attention on visual responses. Vision Res. 49, 1129–1143.

Brainard, D.H. (1997). The Psychophysics Toolbox. Spat. Vis. 10, 433–436.

Briggs, F., Mangun, G.R., and Usrey, W.M. (2013). Attention enhances synaptic efficacy and the signal-to-noise ratio in neural circuits. Nature 499, 476–480.

Brunel, N., and Wang, X.-J. (2001). Effects of Neuromodulation in a Cortical Network Model of Object Working Memory Dominated by Recurrent Inhibition. J. Comput. Neurosci. 11, 63–85.

Buffalo, E.A., Fries, P., Landman, R., Buschman, T.J., and Desimone, R. (2011). Laminar differences in gamma and alpha coherence in the ventral stream. Proc. Natl. Acad. Sci. 108, 11262–11267.

Buia, C.I., and Tiesinga, P.H. (2008). Role of Interneuron Diversity in the Cortical Microcircuit for Attention. J. Neurophysiol. 99, 2158–2182.

Buschman, T.J., and Miller, E.K. (2007). Top-Down Versus Bottom-Up Control of Attention in the Prefrontal and Posterior Parietal Cortices. Science 315, 1860–1862.

Cardin, J.A., Carlén, M., Meletis, K., Knoblich, U., Zhang, F., Deisseroth, K., Tsai, L.-H., and Moore, C.I. (2009). Driving fast-spiking cells induces gamma rhythm and controls sensory responses. Nature 459, 663–667.

Carrasco, M. (2011). Visual attention: The past 25 years. Vision Res. 51, 1484–1525.

Cohen, M.R., and Kohn, A. (2011). Measuring and interpreting neuronal correlations. Nat. Neurosci. 14, 811–819.

Cohen, M.R., and Maunsell, J.H.R. (2009). Attention improves performance primarily by reducing interneuronal correlations. Nat. Neurosci. 12, 1594–1600.

Cohen, M.R., and Maunsell, J.H.R. (2011). Using neuronal populations to study the mechanisms underlying spatial and feature attention. Neuron 70, 1192–1204.

Cowley, B.R., Smith, M.A., Kohn, A., and Yu, B.M. (2016). Stimulus-Driven Population Activity Patterns in Macaque Primary Visual Cortex. PLOS Comput. Biol. 12, e1005185.

Cowley, B.R., Snyder, A.C., Acar, K., Williamson, R.C., Yu, B.M., and Smith, M.A. (2020). Slow Drift of Neural Activity as a Signature of Impulsivity in Macaque Visual and Prefrontal Cortex. Neuron 108, 551–567.e8.

Cunningham, J.P., and Yu, B.M. (2014). Dimensionality reduction for large-scale neural recordings. Nat. Neurosci. 17, 1500–1509.

Dagnino, B., Gariel-Mathis, M.-A., and Roelfsema, P.R. (2014). Microstimulation of area V4 has little effect on spatial attention and on perception of phosphenes evoked in area V1. J. Neurophysiol. 113, 730–739.

Deco, G., and Thiele, A. (2011). Cholinergic control of cortical network interactions enables feedback-mediated attentional modulation. Eur. J. Neurosci. 34, 146–157.

Desimone, R., and Duncan, J. (1995). Neural Mechanisms of Selective Visual Attention. Annu. Rev. Neurosci. 18, 193–222.

Driver, J. (2001). A selective review of selective attention research from the past century. Br. J. Psychol. 92, 53–78.

Ecker, A.S., Denfield, G.H., Bethge, M., and Tolias, A.S. (2016). On the Structure of Neuronal Population Activity under Fluctuations in Attentional State. J. Neurosci. 36, 1775–1789.

Egeth, H.E., and Yantis, S. (1997). VISUAL ATTENTION: Control, Representation, and Time Course. Annu. Rev. Psychol. 48, 269–297.

Elsayed, G.F., and Cunningham, J.P. (2017). Structure in neural population recordings: an expected byproduct of simpler phenomena? Nat. Neurosci. 20, 1310–1318.

Elsayed, G.F., Lara, A.H., Kaufman, M.T., Churchland, M.M., and Cunningham, J.P. (2016). Reorganization between preparatory and movement population responses in motor cortex. Nat. Commun. 7, 13239.

Everitt, B.S. (1984). Maximum Likelihood Estimation of the Parameters in a Mixture of Two Univariate Normal Distributions; a Comparison of Different Algorithms. J. R. Stat. Soc. Ser. Stat. 33, 205–215.

Fries, P. (2015). Rhythms for Cognition: Communication through Coherence. Neuron 88, 220–235.

Fries, W. (1984). Cortical projections to the superior colliculus in the macaque monkey: A retrograde study using horseradish peroxidase. J. Comp. Neurol. 230, 55–76.

Fries, W. (1985). Inputs from motor and premotor cortex to the superior colliculus of the macaque monkey. Behav. Brain Res. 18, 95–105.

Fries, P., Reynolds, J.H., Rorie, A.E., and Desimone, R. (2001). Modulation of oscillatory neuronal synchronization by selective visual attention. Science 291, 1560–1563.

Fu, Y., Tucciarone, J.M., Espinosa, J.S., Sheng, N., Darcy, D.P., Nicoll, R.A., Huang, Z.J., and Stryker, M.P. (2014). A Cortical Circuit for Gain Control by Behavioral State. Cell 156, 1139–1152.

Gilbert, C.D., and Sigman, M. (2007). Brain States: Top-Down Influences in Sensory Processing. Neuron 54, 677–696.

Goldberg, M.E., and Wurtz, R.H. (1972). Activity of superior colliculus in behaving monkey. II. Effect of attention on neuronal responses. J. Neurophysiol. 35, 560–574.

Golub, M.D., Chase, S.M., Batista, A.P., and Yu, B.M. (2016). Brain–computer interfaces for dissecting cognitive processes underlying sensorimotor control. Curr. Opin. Neurobiol. 37, 53–58.

Gregoriou, G.G., Gotts, S.J., Zhou, H., and Desimone, R. (2009). High-frequency, long-range coupling between prefrontal and visual cortex during attention. Science 324, 1207–1210.

Gregoriou, G.G., Rossi, A.F., Ungerleider, L.G., and Desimone, R. (2014). Lesions of prefrontal cortex reduce attentional modulation of neuronal responses and synchrony in V4. Nat. Neurosci. 17, 1003–1011.

Herrero, J.L., Roberts, M.J., Delicato, L.S., Gieselmann, M.A., Dayan, P., and Thiele, A. (2008). Acetylcholine contributes through muscarinic receptors to attentional modulation in V1. Nature 454, 1110–1114.

Herrero, J.L., Gieselmann, M.A., Sanayei, M., and Thiele, A. (2013). Attention-Induced Variance and Noise Correlation Reduction in Macaque V1 Is Mediated by NMDA Receptors. Neuron 78, 729–739.

Huang, C., Ruff, D.A., Pyle, R., Rosenbaum, R., Cohen, M.R., and Doiron, B. (2019). Circuit Models of Low-Dimensional Shared Variability in Cortical Networks. Neuron 101, 337–348.e4.

Ignashchenkova, A., Dicke, P.W., Haarmeier, T., and Thier, P. (2004). Neuron-specific contribution of the superior colliculus to overt and covert shifts of attention. Nat. Neurosci. 7, 56–64.

Indovina, I., and Macaluso, E. (2004). Occipital–parietal interactions during shifts of exogenous visuospatial attention: trial-dependent changes of effective connectivity. Magn. Reson. Imaging 22, 1477–1486.

Jazayeri, M., and Afraz, A. (2017). Navigating the Neural Space in Search of the Neural Code. Neuron 93, 1003–1014.

Kanashiro, T., Ocker, G.K., Cohen, M.R., and Doiron, B. (2017). Attentional modulation of neuronal variability in circuit models of cortex. ELife 6, e23978.

Kanitscheider, I., Coen-Cagli, R., and Pouget, A. (2015). Origin of information-limiting noise correlations. Proc. Natl. Acad. Sci. 112, E6973–E6982.

Karnani, M.M., Jackson, J., Ayzenshtat, I., Sichani, A.H., Manoocheri, K., Kim, S., and Yuste, R. (2016). Opening Holes in the Blanket of Inhibition: Localized Lateral Disinhibition by VIP Interneurons. J. Neurosci. 36, 3471–3480.

Kaufman, M.T., Churchland, M.M., Ryu, S.I., and Shenoy, K.V. (2014). Cortical activity in the null space: permitting preparation without movement. Nat. Neurosci. 17, 440–448.

Kiani, R., Esteky, H., Mirpour, K., and Tanaka, K. (2007). Object Category Structure in Response Patterns of Neuronal Population in Monkey Inferior Temporal Cortex. J. Neurophysiol. 97, 4296–4309.

Klink, P.C., Dagnino, B., Gariel-Mathis, M.-A., and Roelfsema, P.R. (2017). Distinct Feedforward and Feedback Effects of Microstimulation in Visual Cortex Reveal Neural Mechanisms of Texture Segregation. Neuron 95, 209–220.e3.

Kohn, A., Coen-Cagli, R., Kanitscheider, I., and Pouget, A. (2016a). Correlations and neuronal population information. Annu. Rev. Neurosci. 39, 237–256.

Kohn, A., Coen-Cagli, R., Kanitscheider, I., and Pouget, A. (2016b). Correlations and Neuronal Population Information. Annu. Rev. Neurosci. 39, 237–256.

Krauzlis, R.J., Lovejoy, L.P., and Zénon, A. (2013). Superior Colliculus and Visual Spatial Attention. Annu. Rev. Neurosci. 36, 165–182.

Kuchibhotla, K.V., Gill, J.V., Lindsay, G.W., Papadoyannis, E.S., Field, R.E., Sten, T.A.H., Miller, K.D., and Froemke, R.C. (2017). Parallel processing by cortical inhibition enables context-dependent behavior. Nat. Neurosci. 20, 62–71.

Lakatos, P., Karmos, G., Mehta, A.D., Ulbert, I., and Schroeder, C.E. (2008). Entrainment of Neuronal Oscillations as a Mechanism of Attentional Selection. Science 320, 110–113.

Lavie, N. (2010). Attention, Distraction, and Cognitive Control Under Load. Curr. Dir. Psychol. Sci. 19, 143–148.

Lock, T.M., Baizer, J.S., and Bender, D.B. (2003). Distribution of corticotectal cells in macaque. Exp. Brain Res. 151, 455–470.

Luo, T.Z., and Maunsell, J.H.R. (2015). Neuronal Modulations in Visual Cortex Are Associated with Only One of Multiple Components of Attention. Neuron 86, 1182–1188.

Lyon, D.C., Nassi, J.J., and Callaway, E.M. (2010). A Disynaptic Relay from Superior Colliculus to Dorsal Stream Visual Cortex in Macaque Monkey. Neuron 65, 270–279.

Machens, C.K., Romo, R., and Brody, C.D. (2005). Flexible Control of Mutual Inhibition: A Neural Model of Two-Interval Discrimination. Science 307, 1121–1124.

Maunsell, J.H.R. (2015). Neuronal Mechanisms of Visual Attention. Annu. Rev. Vis. Sci. 1, 373–391.

Mayo, J.P., and Maunsell, J.H.R. (2016). Graded Neuronal Modulations Related to Visual Spatial Attention. J. Neurosci. 36, 5353–5361.

Miller, E.K., and Buschman, T.J. (2013). Cortical circuits for the control of attention. Curr. Opin. Neurobiol. 23, 216–222.

Miri, A., Warriner, C.L., Seely, J.S., Elsayed, G.F., Cunningham, J.P., Churchland, M.M., and Jessell, T.M. (2017). Behaviorally Selective Engagement of Short-Latency Effector Pathways by Motor Cortex. Neuron 95, 683–696.e11.

Mitchell, J.F., Sundberg, K.A., and Reynolds, J.H. (2007). Differential Attention-Dependent Response Modulation across Cell Classes in Macaque Visual Area V4. Neuron 55, 131–141.

Mitchell, J.F., Sundberg, K.A., and Reynolds, J.H. (2009). Spatial attention decorrelates intrinsic activity fluctuations in macaque area V4. Neuron 63, 879–888.

Moore, T., and Armstrong, K.M. (2003). Selective gating of visual signals by microstimulation of frontal cortex. Nature 421, 370–373.

Moore, T., and Zirnsak, M. (2017). Neural Mechanisms of Selective Visual Attention. Annu. Rev. Psychol. 68, 47–72.

Morcos, A.S., and Harvey, C.D. (2016). History-dependent variability in population dynamics during evidence accumulation in cortex. Nat. Neurosci. 19, 1672–1681.

Moreno-Bote, R., Beck, J., Kanitscheider, I., Pitkow, X., Latham, P., and Pouget, A. (2014). Information-limiting correlations. Nat. Neurosci. 17, 1410–1417.

Nandy, A.S., Nassi, J.J., and Reynolds, J.H. (2017). Laminar Organization of Attentional Modulation in Macaque Visual Area V4. Neuron 93, 235–246.

Navalpakkam, V., and Itti, L. (2005). Modeling the influence of task on attention. Vision Res. 45, 205–231.

Ni, A.M., Ruff, D.A., Alberts, J.J., Symmonds, J., and Cohen, M.R. (2018). Learning and attention reveal a general relationship between population activity and behavior. Science 359, 463–465.

Nienborg, H., and Cumming, B. (2010). Correlations between the activity of sensory neurons and behavior: how much do they tell us about a neuron’s causality? Curr. Opin. Neurobiol. 20, 376–381.

Nienborg, H. R. Cohen, M., and Cumming, B.G. (2012). Decision-Related Activity in Sensory Neurons: Correlations Among Neurons and with Behavior. Annu. Rev. Neurosci. 35, 463–483.

Oemisch, M., Westendorff, S., Everling, S., and Womelsdorf, T. (2015). Interareal Spike-Train Correlations of Anterior Cingulate and Dorsal Prefrontal Cortex during Attention Shifts. J. Neurosci. 35, 13076–13089.

Ozaki, T.J. (2011). Frontal-to-Parietal Top-Down Causal Streams along the Dorsal Attention Network Exclusively Mediate Voluntary Orienting of Attention. PLOS ONE 6, e20079.

Pandarinath, C., O’Shea, D.J., Collins, J., Jozefowicz, R., Stavisky, S.D., Kao, J.C., Trautmann, E.M., Kaufman, M.T., Ryu, S.I., Hochberg, L.R., et al. (2018). Inferring single-trial neural population dynamics using sequential auto-encoders. Nat. Methods 15, 805–815.

Parker, A.J., and Newsome, W.T. (1998). SENSE AND THE SINGLE NEURON: Probing the Physiology of Perception. Annu. Rev. Neurosci. 21, 227–277.

Peelen, M.V., and Kastner, S. (2014). Attention in the real world: toward understanding its neural basis. Trends Cogn. Sci. 18, 242–250.

Pitkow, X., and Angelaki, D.E. (2017). Inference in the Brain: Statistics Flowing in Redundant Population Codes. Neuron 94, 943–953.

Pooresmaeili, A., Poort, J., and Roelfsema, P.R. (2014). Simultaneous selection by object-based attention in visual and frontal cortex. Proc. Natl. Acad. Sci. 111, 6467–6472.

Recanzone, G.H., and Wurtz, R.H. (2000). Effects of Attention on MT and MST Neuronal Activity During Pursuit Initiation. J. Neurophysiol. 83, 777–790.

Reynolds, J.H., and Chelazzi, L. (2004). Attentional Modulation of Visual Processing. Annu. Rev. Neurosci. 27, 611–647.

Reynolds, J.H., and Heeger, D.J. (2009). The Normalization Model of Attention. Neuron 61, 168–185.

Roberts, M.J., Zinke, W., Guo, K., Robertson, R., McDonald, J.S., and Thiele, A. (2005). Acetylcholine Dynamically Controls Spatial Integration in Marmoset Primary Visual Cortex. J. Neurophysiol. 93, 2062–2072.

Rodman, H.R., Gross, C.G., and Albright, T.D. (1990). Afferent basis of visual response properties in area MT of the macaque. II. Effects of superior colliculus removal. J. Neurosci. 10, 1154–1164.

Rossi, S., Huang, S., Furtak, S.C., Belliveau, J.W., and Ahveninen, J. (2014). Functional connectivity of dorsal and ventral frontoparietal seed regions during auditory orienting. Brain Res. 1583, 159–168.

Rubin, D.B., Van Hooser, S.D., and Miller, K.D. (2015). The Stabilized Supralinear Network: A Unifying Circuit Motif Underlying Multi-Input Integration in Sensory Cortex. Neuron 85, 402–417.

Ruff, D.A., and Cohen, M.R. (2014a). Attention can either increase or decrease spike count correlations in visual cortex. Nat. Neurosci. 17, 1591–1597.

Ruff, D.A., and Cohen, M.R. (2014b). Global cognitive factors modulate correlated response variability between V4 neurons. J. Neurosci. 34, 16408–16416.

Ruff, D.A., and Cohen, M.R. (2016a). Attention Increases Spike Count Correlations between Visual Cortical Areas. J. Neurosci. 36, 7523–7534.

Ruff, D.A., and Cohen, M.R. (2016b). Stimulus Dependence of Correlated Variability across Cortical Areas. J. Neurosci. 36, 7546–7556.

Ruff, D.A., and Cohen, M.R. (2017). A normalization model suggests that attention changes the weighting of inputs between visual areas. Proc. Natl. Acad. Sci. 114, E4085–E4094.

Ruff, D.A., and Cohen, M.R. (2019). Simultaneous multi-area recordings suggest that attention improves performance by reshaping stimulus representations. Nat. Neurosci. 22, 1669–1676.

Ruff, D.A., Alberts, J.J., and Cohen, M.R. (2016). Relating normalization to neuronal populations across cortical areas. J. Neurophysiol. 116, 1375–1386.

Ruff, D.A., Ni, A.M., and Cohen, M.R. (2018). Cognition as a Window into Neuronal Population Space. Annu. Rev. Neurosci. 41, 77–97.

Ruff, D.A., Xue, C., Kramer, L.E., Baqai, F., and Cohen, M.R. (2020). Low rank mechanisms underlying flexible visual representations. Proc. Natl. Acad. Sci. 117, 29321–29329.

Saalmann, Y.B., Pigarev, I.N., and Vidyasagar, T.R. (2007). Neural Mechanisms of Visual Attention: How Top-Down Feedback Highlights Relevant Locations. Science 316, 1612–1615.

Sadtler, P.T., Quick, K.M., Golub, M.D., Chase, S.M., Ryu, S.I., Tyler-Kabara, E.C., Yu, B.M., and Batista, A.P. (2014). Neural constraints on learning. Nature 512, 423–426.

Salinas, E., and Sejnowski, T.J. (2001). Correlated neuronal activity and the flow of neural information. Nat. Rev. Neurosci. 2, 539–550.

Saproo, S., and Serences, J.T. (2014). Attention Improves Transfer of Motion Information between V1 and MT. J. Neurosci. 34, 3586–3596.

Seidemann, E., and Newsome, W.T. (1999). Effect of Spatial Attention on the Responses of Area MT Neurons. J. Neurophysiol. 81, 1783–1794.

Semedo, J., Zandvakili, A., Kohn, A., Machens, C.K., and Yu, B.M. (2014). Extracting Latent Structure From Multiple Interacting Neural Populations. Adv. Neural Inf. Process. Syst. 27.

Semedo, J.D., Zandvakili, A., Machens, C.K., Yu, B.M., and Kohn, A. (2019). Cortical Areas Interact through a Communication Subspace. Neuron 102, 249–259.e4.

Semedo, J.D., Gokcen, E., Machens, C.K., Kohn, A., and Yu, B.M. (2020). Statistical methods for dissecting interactions between brain areas. Curr. Opin. Neurobiol. 65, 59–69.

Semedo, J.D., Jasper, A.I., Zandvakili, A., Aschner, A., Machens, C.K., Kohn, A., and Yu, B.M. (2021). Feedforward and feedback interactions between visual cortical areas use different population activity patterns. BioRxiv 2021.02.08.430346.

Silver, R.A. (2010). Neuronal arithmetic. Nat. Rev. Neurosci. 11, 474–489.

Stepniewska, I., Qi, H.-X., and Kaas, J.H. (1999). Do superior colliculus projection zones in the inferior pulvinar project to MT in primates? Eur. J. Neurosci. 11, 469–480.

Sutherland, M.R., McQuiggan, D.A., Ryan, J.D., and Mather, M. (2017). Perceptual salience does not influence emotional arousal’s impairing effects on top-down attention. Emotion 17, 700–706.

Umakantha, A., Morina, R., Cowley, B.R., Snyder, A.C., Smith, M.A., and Yu, B.M. (2020). Bridging neuronal correlations and dimensionality reduction. BioRxiv 2020.12.04.383604.

Veit, J., Hakim, R., Jadi, M.P., Sejnowski, T.J., and Adesnik, H. (2017). Cortical gamma band synchronization through somatostatin interneurons. Nat. Neurosci. 20, 951–959.

Verhoef, B.-E., and Maunsell, J.H.R. (2017). Attention-related changes in correlated neuronal activity arise from normalization mechanisms. Nat. Neurosci. 20, 969–977.

Womelsdorf, T., and Fries, P. (2007). The role of neuronal synchronization in selective attention. Curr. Opin. Neurobiol. 17, 154–160.

Womelsdorf, T., Fries, P., Mitra, P.P., and Desimone, R. (2006a). Gamma-band synchronization in visual cortex predicts speed of change detection. Nature 439, 733–736.

Womelsdorf, T., Anton-Erxleben, K., Pieper, F., and Treue, S. (2006b). Dynamic shifts of visual receptive fields in cortical area MT by spatial attention. Nat. Neurosci. 9, 1156–1160.

Yan, Y., Rasch, M.J., Chen, M., Xiang, X., Huang, M., Wu, S., and Li, W. (2014). Perceptual training continuously refines neuronal population codes in primary visual cortex. Nat. Neurosci. 17, 1380–1387.

Yu, B.M., Cunningham, J.P., Santhanam, G., Ryu, S.I., Shenoy, K.V., and Sahani, M. (2009). Gaussian-Process Factor Analysis for Low-Dimensional Single-Trial Analysis of Neural Population Activity. J. Neurophysiol. 102, 614–635.

Zénon, A., and Krauzlis, R.J. (2012). Attention deficits without cortical neuronal deficits. Nature 489, 434–437.

